# Ras-dependent RAF-MAPK hyperactivation by pathogenic RIT1 is a therapeutic target in Noonan syndrome-associated cardiac hypertrophy

**DOI:** 10.1101/2022.11.02.514888

**Authors:** Antonio Cuevas-Navarro, Morgan Wagner, Richard Van, Monalisa Swain, Madeline R. Allison, Alice Cheng, Simon Messing, Dhirendra K. Simanshu, Matthew J. Sale, Frank McCormick, Andrew G. Stephen, Pau Castel

## Abstract

RIT1 belongs to the family of Ras guanosine triphosphatases (GTPases) that regulate many aspects of signal transduction and are drivers of cancer and congenital disorders. *RIT1* gain-of-function mutations are found in lung cancer, leukemia, and in the germline of Noonan syndrome individuals with an increased prevalence of cardiac hypertrophy and other congenital heart defects. Pathogenic RIT1 proteins evade proteasomal degradation and promote MEK/ERK mitogen-activated protein kinase (MAPK) hyperactivation, yet the mechanism remains poorly understood. Here we show that RAF kinases are putative mutant RIT1 effectors necessary for MAPK activation and characterize RIT1 association with plasma membrane lipids and interaction with RAF kinases. We identify critical residues present in the RIT1 hypervariable region that facilitate interaction with negatively charged membrane lipids and show that these are necessary for association with RAF kinases. Although mutant RIT1 binds to RAF kinases directly, it fails to activate RAF-MAPK signaling in the absence of classical Ras proteins. Consistent with aberrant RAF/MEK/ERK activation as a driver of disease, we show that MEK inhibition alleviates cardiac hypertrophy in a mouse model of RIT1-mutant Noonan syndrome. These data shed light on pathogenic RIT1 function and identify avenues for therapeutic intervention.

**One Sentence Summary:** Electrostatic plasma membrane association facilitates RIT1-mediated Ras-dependent RAF kinase activation to promote pathogenic MAPK signaling.

## INTRODUCTION

The Ras family of small guanosine triphosphatases (GTPases) control a diverse network of signaling pathways essential for human development and adult tissue homeostasis (*1, 2*). The activity of Ras GTPases is dictated by their nucleotide-loading state and is subject to various regulatory mechanisms that respond to extracellular stimuli and feedback signals (*2*). Ras GTPases bound to guanosine diphosphate (GDP) are considered inactive, while their guanosine triphosphate (GTP)-bound conformation promotes activation of downstream effectors. GTPase activating proteins (GAPs), such as NF1 and RASA1/p120GAP, turn off Ras proteins by catalyzing GTP hydrolysis. Conversely, guanine nucleotide exchange factors (GEFs), such as SOS1/2, activate Ras by promoting the exchange of GDP for GTP. When bound to GTP, the classical Ras proteins HRAS, NRAS, and KRAS (hereinafter referred to as Ras) recruit RAF kinases and other effector proteins to the plasma membrane (PM) to activate various downstream signaling cascades. Indeed, translocation of RAF to the PM, mediated by its high-affinity Ras-binding domain (RBD), is a crucial step in the activation of the RAF/MEK/ERK MAPK pathway (*3, 4*). Hyperactivation of this critical Ras signaling axis is a hallmark of many cancers and the cause of a group of developmental syndromes collectively called RASopathies (*5*).

Recently, gain-of-function mutations in the Ras-related GTPase RIT1 have emerged as drivers of human disease, including cancers of the lung and myeloid malignancies (*6*–*8*). Germline *RIT1* variants cause Noonan syndrome (NS), a RASopathy characterized by craniofacial dysmorphism, short stature, and congenital heart disease (*9*–*11*). RIT1 NS patients make up approximately 5-10% of all NS cases and exhibit congenital cardiac defects at elevated frequencies (*9, 12*). Recent murine models of RIT1 NS, independently developed by our group and others, present features that recapitulate clinical manifestations, including a shortened stature, craniofacial dysmorphism, and cardiac hypertrophy (*13, 14*), providing an avenue for the evaluation of therapies against hypertrophic cardiomyopathy (HCM) and other cardiac defects closely associated with RIT1 NS in a preclinical setting.

RIT1 is expressed in many tissues and, like other Ras GTPases, associates with the inner leaflet of the PM through a unique C-terminal hypervariable region (HVR). The G-domain of RIT1 shares 51% sequence identity with Ras and thus can potentially associate with an overlapping set of effector proteins, including the RAF kinases. However, regulation of the RIT1 GTPase cycle remains elusive, with cognate RIT1 GAPs and GEFs yet to be identified. Interestingly, a high fraction of cellular RIT1 is bound to GTP, even in the absence of mitogenic signals, suggesting that RIT1 activity may rely on its intrinsic nucleotide exchange and hydrolase activity and/or alternative regulatory mechanisms (*13*). One such mechanism involves the NS-associated protein LZTR1, which functions as a conserved substrate receptor for Cullin3 E3 ubiquitin ligase complexes (CRL3^LZTR1^) to promote the ubiquitination and proteasomal degradation of GDP-bound RIT1 (*15*). NS pathogenic RIT1 and LZTR1 variants disrupt RIT1-LZTR1 binding, resulting in the accumulation of RIT1 protein (*13, 16*) and enhanced MAPK signaling; however, it remains unclear how increased RIT1 protein abundance contributes to MAPK pathway activation.

Here, we employ biophysical, biochemical, and cell biological approaches to interrogate RIT1 activation of RAF kinases. We show that different biochemical properties of the RIT1 HVR contribute to its association with the PM and enable RAF binding. RIT1 exhibits preferential binding to RAF1 (also known as CRAF) and engages with an overlapping set of RBD residues associated with Ras-binding, albeit with a weaker affinity. Furthermore, we demonstrate that the absence of Ras proteins limits the ability of pathogenic RIT1 to hyperactivate the MAPK pathway and that pharmacological MAPK inhibition ameliorates cardiac tissue overgrowth associated with aberrant RIT1 signaling in a RIT1 NS mouse model.

## RESULTS

### RIT1 oncoproteins activate RAF kinase

Although expression of pathogenic RIT1 variants has been widely demonstrated to promote MEK/ERK pathway activation, the mechanism by which RIT1 activates MAPK signaling remains unclear (*6, 10, 11, 13*). RIT1 associates with RAF kinases in a nucleotide-dependent manner (*17*), a defining feature of GTPase effector proteins, postulating a direct link between RIT1 and MEK/ERK activation. To directly assess whether RIT1 oncoproteins activate MAPK signaling in a RAF-dependent manner, we expressed mutant RIT1 and assessed ERK phosphorylation following treatment with a third-generation small molecule pan-RAF inhibitor (*18*). ERK activation induced by expression of RIT1^A57G^, RIT1^M90I^, and RIT1^T83P^, three pathogenic variants with variable levels of GTP-loading in HEK293T cells (*13*), was abrogated by pharmacological RAF inhibition (**Fig. 1A**). Furthermore, knockdown of the three RAF isoforms (ARAF, BRAF, and RAF1) by RNAi reduced mutant RIT1-induced ERK phosphorylation, highlighting a dependency on RAF kinases (**Fig. 1B**). To determine whether mutant RIT1 expression activates RAF proteins, we measured the in vitro kinase activity of RAF isolated from mammalian cells co-expressing RIT1^A57G^, the most common RIT1 NS variant, or an empty vector (EV) control, cultured in the absence of serum (**Fig. 1C and fig. S1**). As previously noted, BRAF exhibited high basal activity in vitro (*19*), potentially due to constitutive phosphorylation of its activation loop (*20*), but was largely refractory to RIT1^A57G^ co-expression. In contrast, the kinase activity of RAF1 was significantly enhanced by RIT1^A57G^ co-expression. Together, these data suggest that mutant RIT1 activates RAF kinases in cells and that the RAF-MEK-ERK cascade is a putative effector pathway of mutant RIT1.

**Fig. 1.**
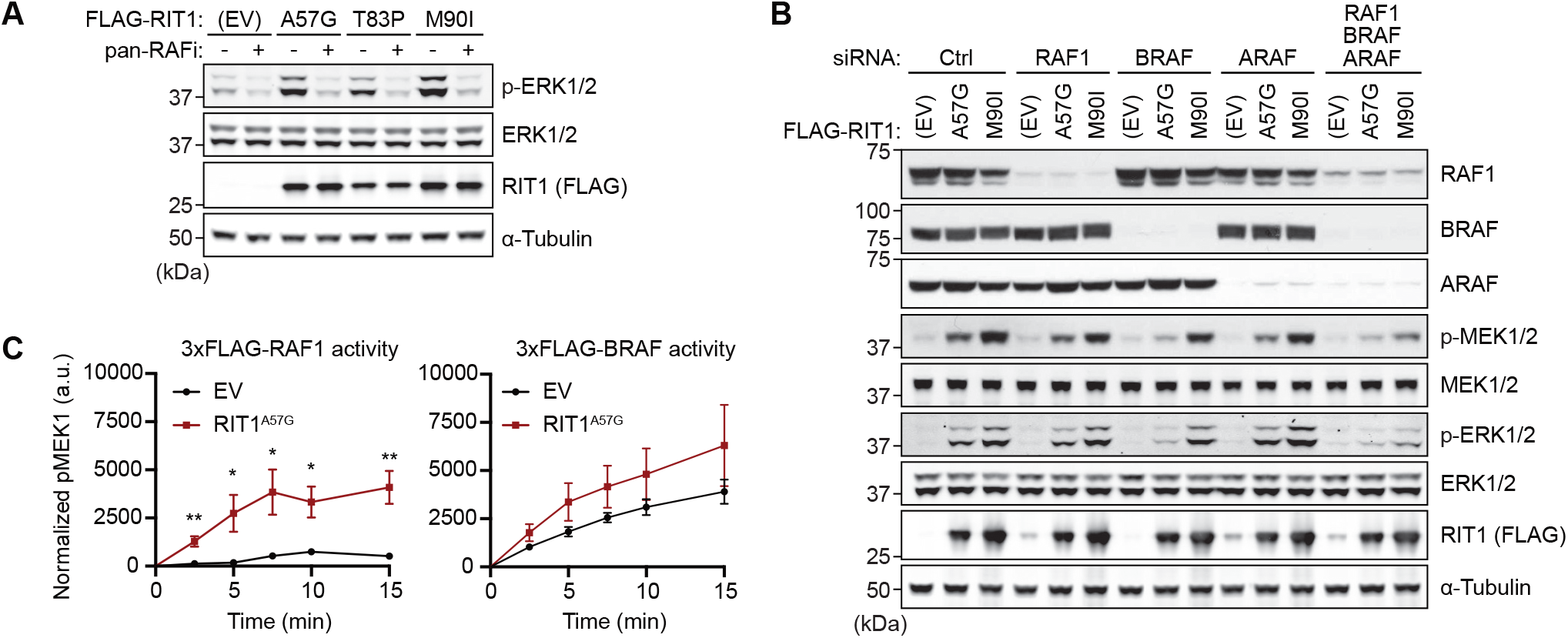
RIT1 oncoproteins activate ERK signaling via RAF kinase. (**A**) Immunoblot analysis of indicated proteins from HEK293T cells transiently transfected with indicated FLAG-tagged RIT1 constructs or an empty vector (EV) control. Cells were serum-starved for 16 h then treated with 10 µM LY3009120 (pan-RAFi) or DMSO vehicle control for 1 h. One of three independent experiments is shown. (**B**) Immunoblot analysis of indicated proteins from HEK293T cells transiently transfected with indicated siRNA and FLAG-tagged RIT1 constructs or EV control. One of two independent experiments is shown. (**C**) in vitro MEK1 phosphorylation (Ser^218^/Ser^222^) by RAF1 or BRAF protein isolated from HEK293T cells co-expressing RIT1^A57G^ or an EV control. a.u., arbitrary units. Data points indicate the mean ± SEM of four (RAF1) or three (BRAF) biological replicates (independently isolated RAF protein samples).

### RIT1 association with the PM requires charge complementarity

Given the crucial role played by the PM in the activation of RAF kinases, we sought to investigate the association of RIT1 with the inner leaflet of the PM. Unlike classical Ras proteins, the RIT1 HVR lacks prenylation motifs, indicating that RIT1 engages with the PM in a unique way. To investigate this interaction, we used surface plasmon resonance (SPR) to measure the association of RIT1 with liposomes of various charge ratios (**Fig. 2A**). RIT1 showed no binding response to neutral liposomes, however as the negative charge in the liposome was increased via the inclusion of 16:0-18:1 phosphatidylserine (POPS), an increasing binding response was observed (**Fig. 2B**). We confirmed this interaction was strictly mediated by the HVR because upon its removal we did not observe interaction with liposomes containing 30% POPS (**fig. S2A**). The association of RIT1 with anionic lipid-containing liposomes was quantitated by calculating the partition coefficient (*21*)(**fig. S2B**). Increasing the concentration of POPS in the liposomes from 10 to 30% increased the partition coefficient by 40-fold. Consistent with our data, molecular dynamic simulations have found that the RIT1 HVR experiences a significantly longer residence time on POPS-containing bilayers compared to uncharged bilayers (*22*).

**Fig. 2.**
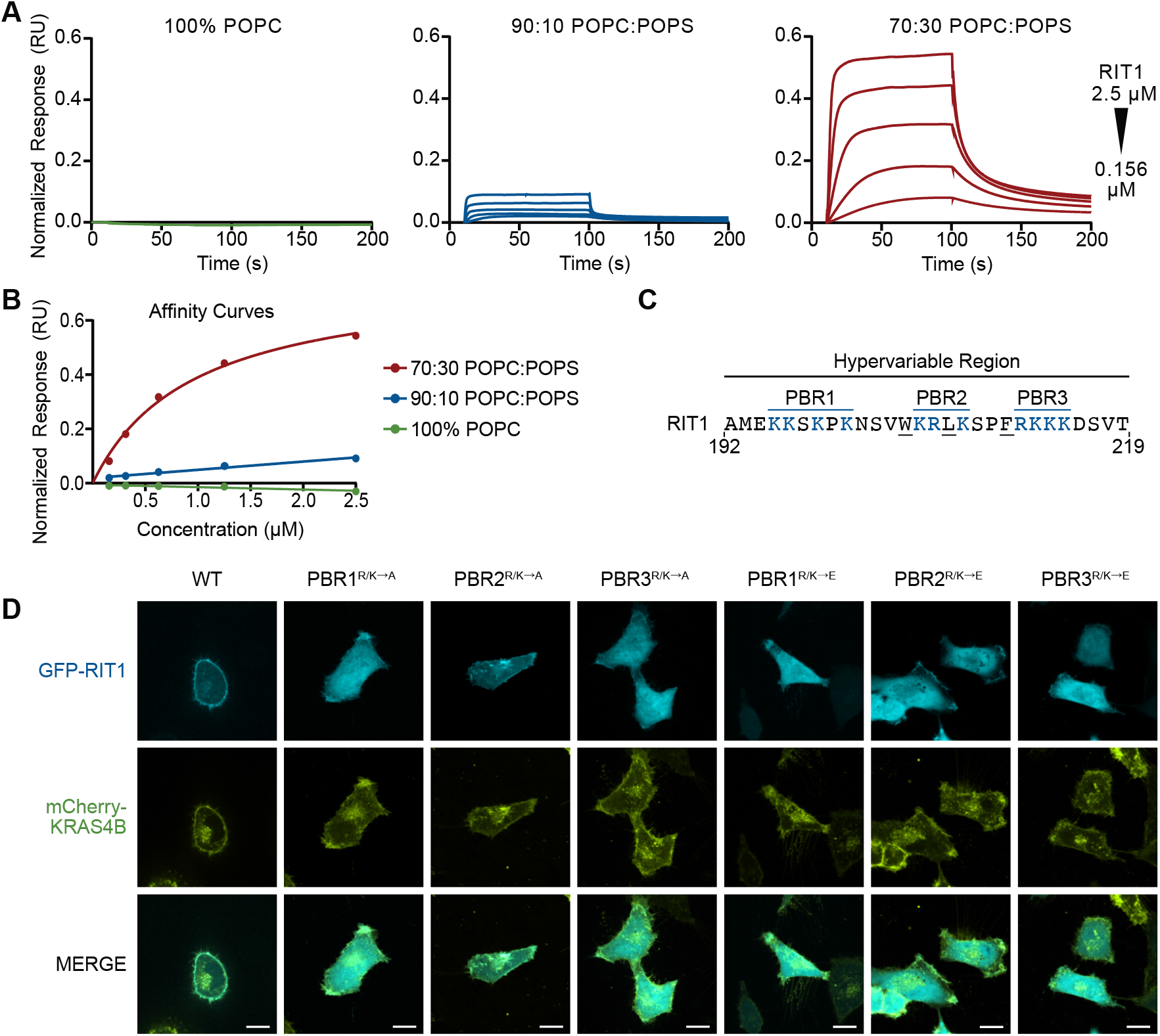
RIT1 associates with the plasma membrane through its HVR. (**A**) SPR analysis with increasing concentrations of POPS-containing liposomes showing RIT1 (a.a. 17-219) association with negatively charged lipids. (**B**) SPR affinity curves showing relative binding affinity of RIT1 to lipid constructs. (**C**) RIT1 C-terminal amino acid sequence. Polybasic regions (PBR) are colored blue. (**D**) Live-cell confocal images of HeLa cells transiently transfected with indicated GFP-RIT1 constructs. Stable expression of mCherry-KRAS4B was used as a plasma membrane marker. Representative images from one of three independent experiments (*n* = 3). Scale bar is 15 μm.

The RIT1 HVR has an isoelectric point of 10.6 and contains three polybasic regions (PBR) separated by non-charged amino acids (**Fig 2C**). To better characterize the contribution of these PBRs, we assessed the PM association of green fluorescent protein (GFP)-tagged RIT1 with individual PBRs that had their charge neutralized (R/K→A) or reversed (R/K→E) (**Fig 2D**). As expected, charge reversal and neutralization of PBR1, PBR3, and to a lesser degree PBR2, disrupted the typical distribution of RIT1 at the cellular periphery, suggesting that all three PBRs are essential for membrane association in cells. PBR2 contains fewer basic residues than PBR1 and PBR3 and, thus, the contribution to membrane association provided by each PBR may be directly correlated to their overall charge contribution. In addition, charge neutralization of a single basic residue within PBR2 or PBR3 (Arg^206^ and Arg^212^, respectively) was insufficient to disrupt membrane association (**fig. S2C**). Using a GFP-RIT1 C-terminal peptide fusion construct, Heo *et al*., (2006) demonstrated that hydrophobic side chains of the RIT1 HVR may also regulate RIT1-PM association (*23*). Therefore, we individually mutated five hydrophobic HVR residues in a full-length GFP-RIT1 construct. Of these, alanine substitution of the three largest side chains disrupted PM targeting (**fig. S2C**). Collectively, these data indicate that charge complementarity plays a significant role in the association of RIT1 with the inner leaflet of the PM and that this interaction receives a contribution from the hydrophobic residues interspersed between the PBRs.

### Characterization of RIT1-RAF interactions

The HVRs of classical Ras proteins contribute to distinct binding preferences with the different RAF family members (*24*). Given RIT1’s unique HVR, we sought to determine whether RIT1 also exhibits preferential RAF isoform binding. Notably, binding of the three RAF isoforms to wild-type (WT) RIT1 was nearly undetectable in pulldown assays and was markedly weaker than their affinity to KRAS4B (**Fig. 3A**). However, in contrast to prior observations (*17*), a significant preference for RAF1 was observed when using the pathogenic RIT1^A57G^ variant. Thus, to better assess these interactions, we developed a quantitative bioluminescence resonance energy transfer (BRET) assay to quantitate the association of RIT1 with RAF kinases in cells (**fig. S3A, B**). WT RIT1 had higher BRET_max_ and lower BRET_50_ values for RAF1 compared to BRAF and ARAF, indicating a preference for the RAF1 paralog in intact cells (**Fig. 3B and fig. S3C)**. Based on the calculated BRET_50_ values, RIT1 binds preferentially to RAF1 over BRAF and ARAF. Remarkably, RIT1^A57G^ bound to RAF1 around 10-fold tighter than WT RIT1 and showed the same binding preferences as the WT protein: RAF1>BRAF>ARAF. To investigate this interaction further we used isothermal titration calorimetry (ITC) to measure the binding affinity between recombinant RAF1- or BRAF-RBD and WT RIT1 or RIT1^A57G^ bound to GMPPNP, a non-hydrolyzable GTP analog (**Fig. 3C**). In these experiments, RIT1^A57G^ (*K*_*D*_ = 2.96 µM) bound approximately 5-times more tightly to RAF1-RBD compared with WT (*K*_*D*_ = 14.55 µM). Consistent with our BRET measurements, RIT1^A57G^ interaction with BRAF-RBD was weaker than RAF1-RBD. However, no measurable binding was observed between WT RIT1 and BRAF-RBD, presumably because the binding interaction was below the detection limit for ITC. Cumulatively, these data suggest that RIT1 preferentially interacts with RAF1 over the other RAF isoforms.

**Fig. 3.**
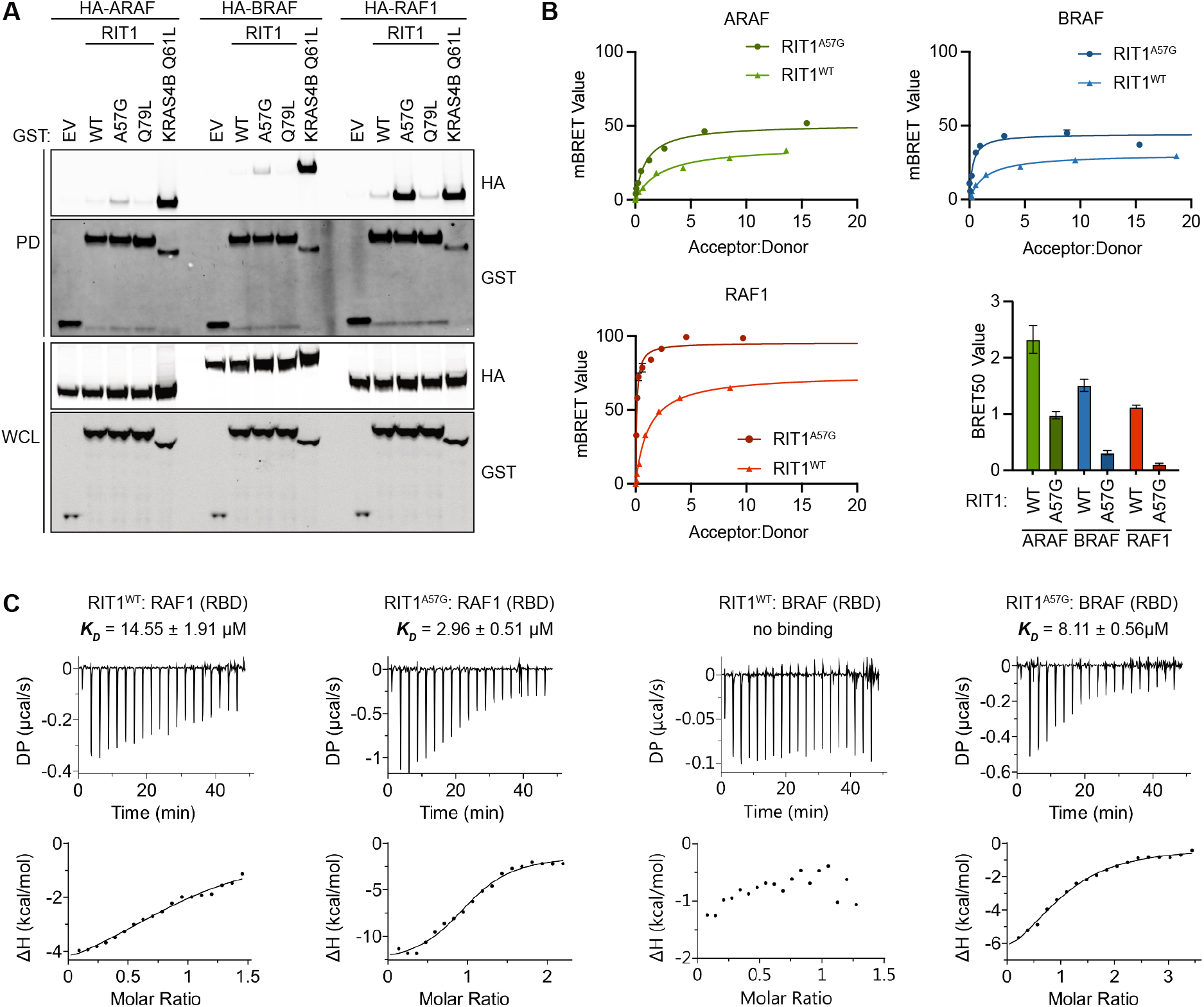
RIT1 exhibits preferential binding to RAF1. (**A**) Immunoblot analysis of indicated proteins precipitated by GST pulldown assay from HEK293T cell lysates expressing indicated constructs. DNA amounts for WT and mutant RIT1 were adjusted to normalize for protein expression. EV, empty vector; WCL, whole-cell lysate. RIT1^Q79L^ is the corresponding mutant of RAS^Q61L^. (**B**) BRET curves show the relative binding affinities of mVenus-RIT1 (acceptor) and RAF-nanoLuc (donor) proteins. Representative BRET curves from three independent experiments are shown. The histogram demonstrates the mean BRET_50_ values ± SD of three independent experiments. (**C**) ITC measurements of recombinant RIT1:RAF(RBD) binding affinities. K_D_ values represent an average of three independent experiments.

To understand how RAF binding differs between RIT1 and RAS, we used solution nuclear magnetic resonance (NMR) spectroscopy to identify broadened and induced chemical shifts upon RBD binding (**Table S1 and fig. S4**). An overlay of the RAF1-RBD chemical shift perturbation (CSP) histograms produced by binding to WT RIT1, RIT1^A57G^, or KRAS revealed an overlapping set of perturbed residues with minor variations (**Fig. 4A**). Of note, differences in binding affinities were evident by CSP analysis and were congruent with affinities measured in vitro and in cells (**Fig. 3**); specifically, WT RIT1 induced the smallest perturbation of resonances, followed by RIT1^A57G^, then KRAS, which induced the largest perturbations. Perturbed residues were then mapped onto modeled structures of WT and A57G mutants of RIT1 in the active state and were compared with the previously solved crystal structure of the KRAS:RAF1-RBD complex (*25*)(**Fig. 4B**). This identified the putative RIT1:RBD interface and confirmed a shared binding site on RAF1-RBD. CSP analysis of RIT1 revealed residues (Met^37^, Ser^43^, and His^44^) at the N-terminal end of the switch-I flexible loop, respectively, that were considerably perturbed in the A57G mutant but not in the WT form (**Fig. 4C, D, Table S1, and fig. S4D-F**). These data suggest that the switch-I of RIT1^A57G^ differentially engages RAF1-RBD, potentially providing the enhanced stability exhibited by the pathogenic variant.

**Fig. 4.**
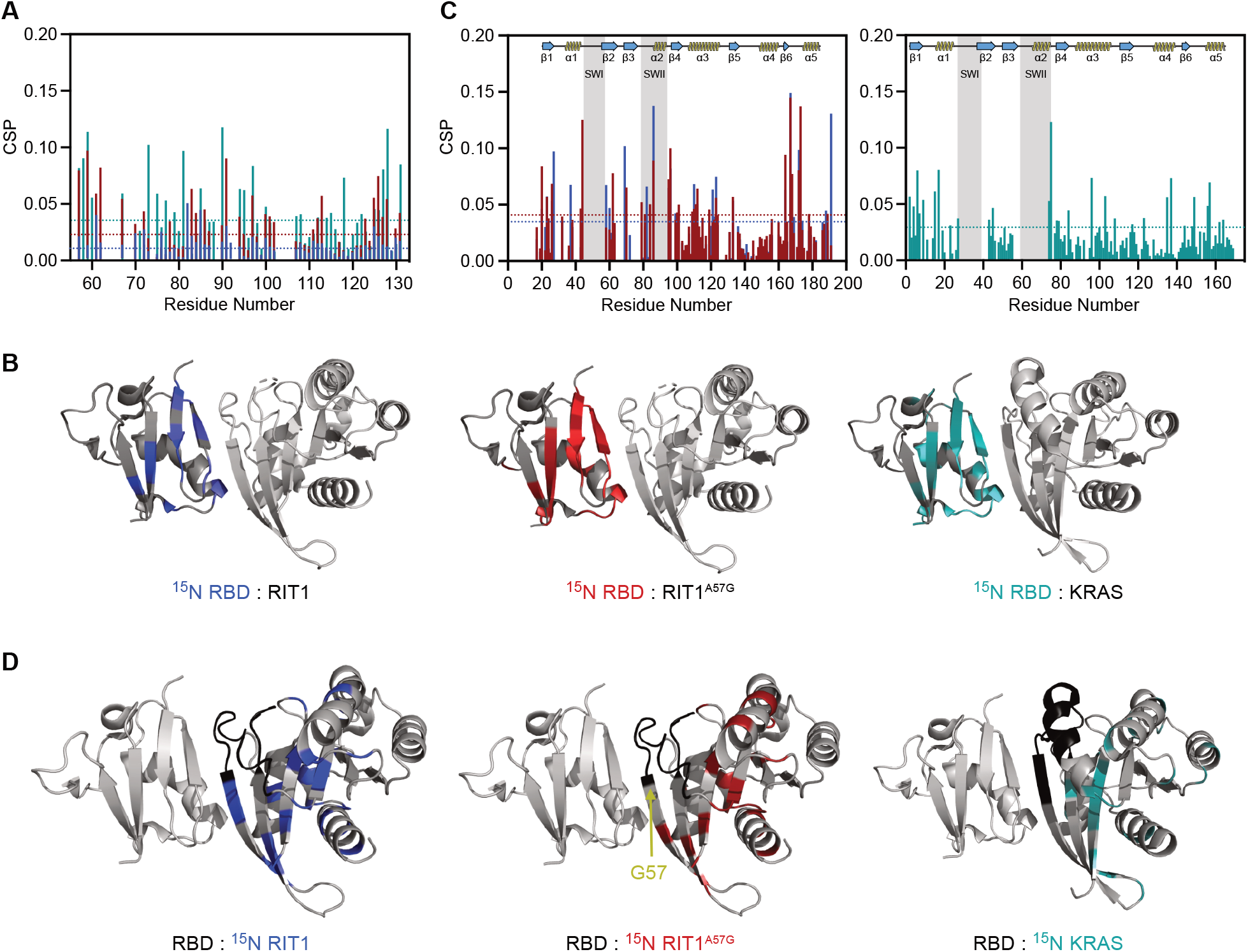
Characterization of RIT1-RAF RBD interface by NMR. (**A**) Chemical shift perturbations (CSP) plots for ^15^N RAF1-RBD observable in complex with unlabeled WT RIT1 (blue), RIT1^A57G^ (red) & WT KRAS (cyan). Dashed lines represent 1.5σ. (**B**) CSP shown in (A) and broadened residues were mapped to the 3D-modelled RBD-RIT1 complex and RBD-KRAS structure (PDB: 6VJJ). (**C**) Left panel represents the CSP for ^15^N WT RIT1 (blue) and ^15^N RIT1^A57G^ (red) in complex with unlabeled RBD. The right panel represents CSP for ^15^N WT KRAS in complex with unlabeled RAF1-RBD. Dashed lines represent 1.5σ. (**D**) CSP shown in (C) and broadened residues were mapped to the 3D structures as in (B).

To further interrogate how the A57G mutation in RIT1 enhances its interaction with RAF1, we undertook a comparative analysis of the modeled RIT1:RAF1-RBD structures with that of the solved KRAS-RAF1-RBD structure (*25*)(**Fig. 5A**). The RIT1 Ala^57^ residue located at the end of the switch-I region is equivalent to the Ser^39^ residue in KRAS (**Fig. 5B**). In the KRAS:RAF1-RBD complex structure, side-chain and main-chain atoms of Ser^39^ form hydrogen (H-) bonds with Arg^67^ and Arg^89^ residues of RAF1. KRAS residues Asp^38^ and Tyr^40^, which surround amino acid Ser^39^, form key interactions with RAF1-RBD by forming salt-bridge, H-bond, and van der Waals interaction with RAF1 Arg^89^ and Thr^68^ residues (*25*). R89L mutation in RAF1 and mutations of Ser^39^ neighboring residues (E37G, D38A, and Y40C mutations) in Ras proteins have been shown to result in either complete or significant loss of binding between KRAS and RAF1-RBD (*21, 25*–*27*), suggesting the interactions formed by Ser^39^ and residues around it in KRAS and Arg^89^ in RAF1 play a critical role in KRAS:RAF interaction. Unlike the KRAS Ser^39^ residue, the Ala^57^ or Gly^57^ (A57G) residues in RIT1 cannot form an H-bond with Arg^67^ as they lack a side-chain hydroxyl group (**Fig. 5B, C**). Glycine is fundamentally different from Ala and all other amino acids in that it lacks a side chain, which allows a much larger rotational freedom of its main-chain torsion angles (*28*). In the beta-strand, glycine is often present at N-cap and C-cap positions, and the presence of glycine in the middle of the beta-strand has been shown to have a strong tendency to block β-sheet continuation (*29*). Thus, unlike Ala^57^ in WT RIT1, the Gly^57^ residue in RIT1 possesses increased rotational space of main-chain torsion angles, imparting flexibility to the local peptide structure. This allows Gly^57^ and neighboring residues in switch-I of RIT1 to undergo minor conformational changes that permit higher-affinity interactions with RAF1-RBD, as suggested by the additional perturbation and broadening of RAF1 interface residues Arg^67^ and Val^70^, respectively, similar to the conformational changes observed upon KRAS:RAF1-RBD complex formation (**Fig. 5B-D, Table S1**). Thus, the A57G mutation likely enhances the RIT1:RAF1 interaction through increased flexibility in its main-chain torsion angles, which impact not only Gly^57^ but also neighboring residues, including switch-I and flanking residues (Ser^43^ and His^44^) (**Fig. 4C**). Together, these play a crucial role in forming an enhanced interaction with RAF1-RBD.

**Fig. 5.**
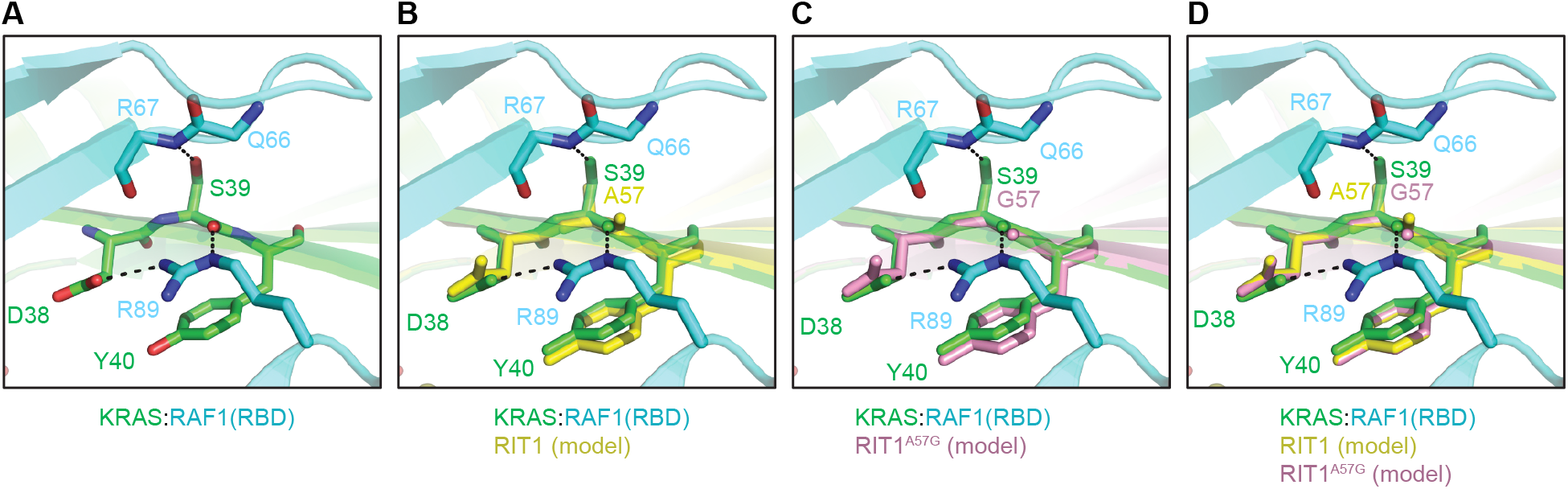
Analysis of modeled RIT1-RAF RBD interface. (**A**-**D**) Comparison of modeled structures of WT and A57G mutant of RIT1 with the crystal structure of KRAS-RAF1(RBD) complex (PDB: 6VJJ). (A) Interaction formed by KRAS Ser^39^ (equivalent to Ala^57^ in RIT1) and neighboring residues Asp^38^ and Tyr^40^ with RAF1-RBD. KRAS and RAF1-RBD are colored green and cyan, respectively. (B) Superposition of KRAS-RAF1(RBD) complex with the modeled structure of WT RIT1 (colored yellow) in the active state. (C) Superposition of KRAS-RAF1(RBD) complex with the modeled structure of A57G mutant of RIT1 (colored pink) in the active state. (D) Superposition of KRAS-RAF1(RBD) complex with the modeled structures of WT and A57G mutant of RIT1 in the active state.

### RIT1 membrane localization is required for RAF interaction

Ras:RAF-RBD binding is insufficient for RAF activation in the absence of vicinal PM phospholipids and thus PM anchoring is a requisite for Ras-driven MAPK activation (*30*). Therefore, we reasoned that RIT1:RAF binding and activation may exhibit similar dependency on PM association. To assess the role of RIT1 membrane trafficking on RAF activation, we expressed N-terminal and C-terminal RIT1 deletion mutants and assessed binding by pulldown. As predicted, deletion of the RIT1 C-terminus (192-219), but not its N-terminus (1-18), disrupted membrane association, RAF1 binding, and MAPK activation (**Fig. 6A and fig. S5A**). Importantly, rescuing membrane association of the RIT1 mutant lacking HVR by introducing a CAAX box motif, restored RIT1:RAF1 binding and activation of MAPK, indicating that PM localization is required for productive RAF interaction. Notably, when measuring relative RIT1 GTP-loading in a RAF1-RBD pulldown assay, deletion of the RIT1 C-terminus had no effect on GTP-dependent RBD binding that was abrogated by a dominant negative S35N mutation (*17, 31*), equivalent to the S17N in Ras which stabilizes GDP-binding (*32*) (**fig. S5B**). These data suggest that, while necessary for RIT1-mediated RAF activation, PM binding is dispensable for GTP loading.

**Fig. 6.**
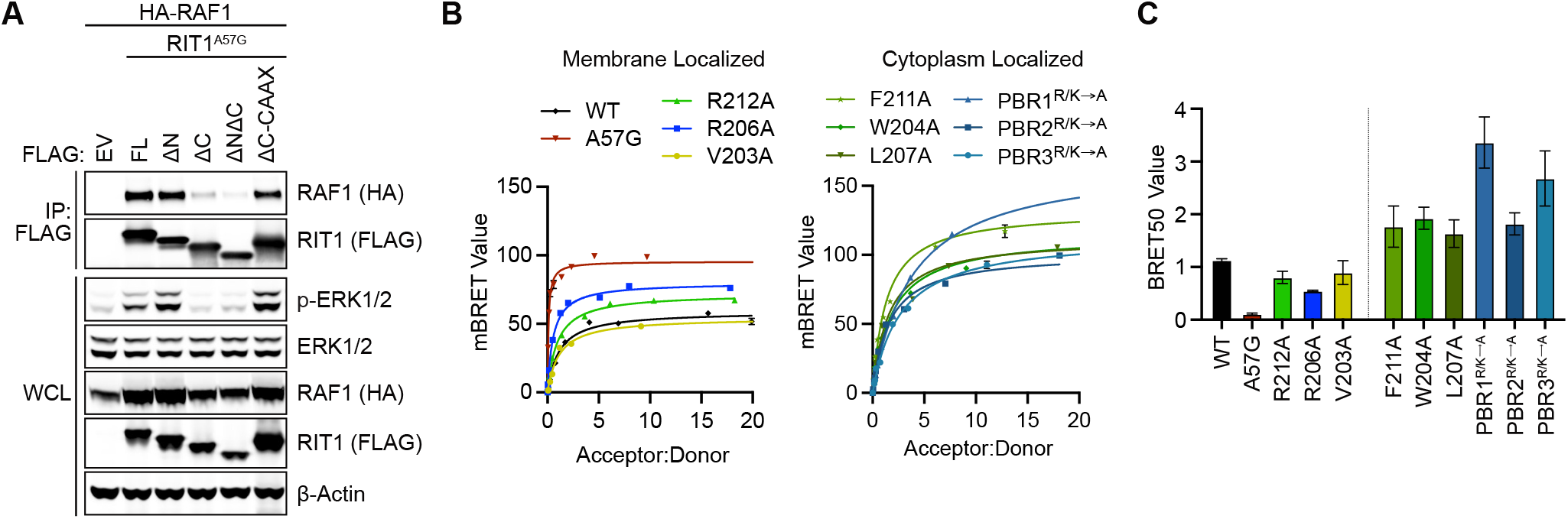
RIT1 HVR is required for RAF binding. (**A**) Immunoblot analysis of indicated proteins immunoprecipitated from HEK293T cell lysates expressing indicated constructs. EV, empty vector; ΔN, a.a. 1-18 deletion; ΔC, a.a. 192-219 deletion; WCL, whole-cell lysate. (**B**) BRET assays of indicated C-terminal HVR mutants associated with RAF1-nanLuc. One of three experiments is shown. (**C**) Histogram of BRET_50_ values indicating the relative binding affinities of RIT1 C-terminal mutants for RAF1-nanoLuc. Mean BRET_50_ values ± SD of three independent experiments.

Given the critical function of the HVR in membrane localization, we sought to quantitatively assess how its biochemical properties contribute to RIT1:RAF1 binding in vivo. Point mutations that had no impact on membrane localization (**fig. S2C**), such as V203A, R206A, and R212A, bound RAF1 with a BRET_50_ comparable to WT RIT1 (**Fig. 6B, C)**. However, mutations that significantly decreased membrane localization produced much larger BRET_50_ values, indicating a weaker association with RAF1. Interestingly, charge neutralizing mutations in PBR 1 and 3 had the largest impact on RAF1 binding, consistent with their effect on RIT1 PM localization (**Fig. 2D**). Together, these findings suggest that while the RIT1 HVR does not directly participate in the activation of RAF, it is necessary for RIT1 localization to the PM where it can interact with RAF and activate the MAPK cascade.

### Mutant RIT1 activates RAF in a Ras-dependent manner

Given that RIT1 binds RAF directly, one can posit that RIT1, like Ras, may promote downstream MAPK signaling via direct recruitment and activation of RAF at the PM and that MAPK activation may be limited by RIT1’s weak affinity towards RAF (*13*). Intriguingly, despite an increased affinity towards RAF, RIT1^A57G^ exhibits comparable MAPK activation relative to other pathogenic variants with an affinity towards RAF that is indistinguishable from WT RIT1, including the oncogenic M90I allele (*13, 33*). This suggests that the activation mechanism may be subject to limiting factors independent of IT1-RAF binding. Given that other non-classical Ras GTPases, such as MRAS (*34, 35*), promote RAF activation in a Ras-dependent manner, we hypothesized that pathogenic RIT1 may also rely on classical Ras proteins to activate RAF. Therefore, to evaluate Ras dependency, we knocked down HRAS, NRAS, and KRAS in WT or RIT1^M90I^ expressing primary mouse embryonic fibroblasts (MEFs). As expected, Ras knockdown attenuated growth-factor-induced MAPK activation (**fig. S6A**). Moreover, RIT1^M90I^ enhanced the ERK signaling response compared to WT and its effect was attenuated by Ras knockdown. However, despite our best efforts to deplete Ras cells using RNAi, this approach’s inefficiency did not allow for a proper evaluation of Ras dependency. Therefore, we generated HEK293 cells devoid of classical Ras proteins via CRISPR/Cas9-mediated triple knockout (TKO) of *HRAS, NRAS*, and *KRAS*, which rendered them insensitive to RTK-mediated MAPK pathway activation (**fig. S6B, C**). The “Rasless” 293 system was then used to evaluate the role of Ras proteins in mutant RIT1-MAPK signaling. Ectopic expression of two pathogenic RIT1 alleles (A57G and M90I) and RIT1^Q79L^ activated MAPK signaling in control cells but not in Rasless 293 TKO cells (**Fig. 7A**). Ectopic rescue of Ras expression in TKO cells reinstated RIT1-mediated MAPK signaling (**Fig. 7A)**, suggesting that in this cell system, RIT1 relies on Ras proteins to activate the MAPK pathway. Furthermore, the addition of a dominant-negative S35N mutation or a C-terminal deletion mutant confirmed that MAPK activation in control cells was dependent on effector binding and proper localization of RIT1 to the PM, respectively (**fig. S6D**).

**Fig. 7.**
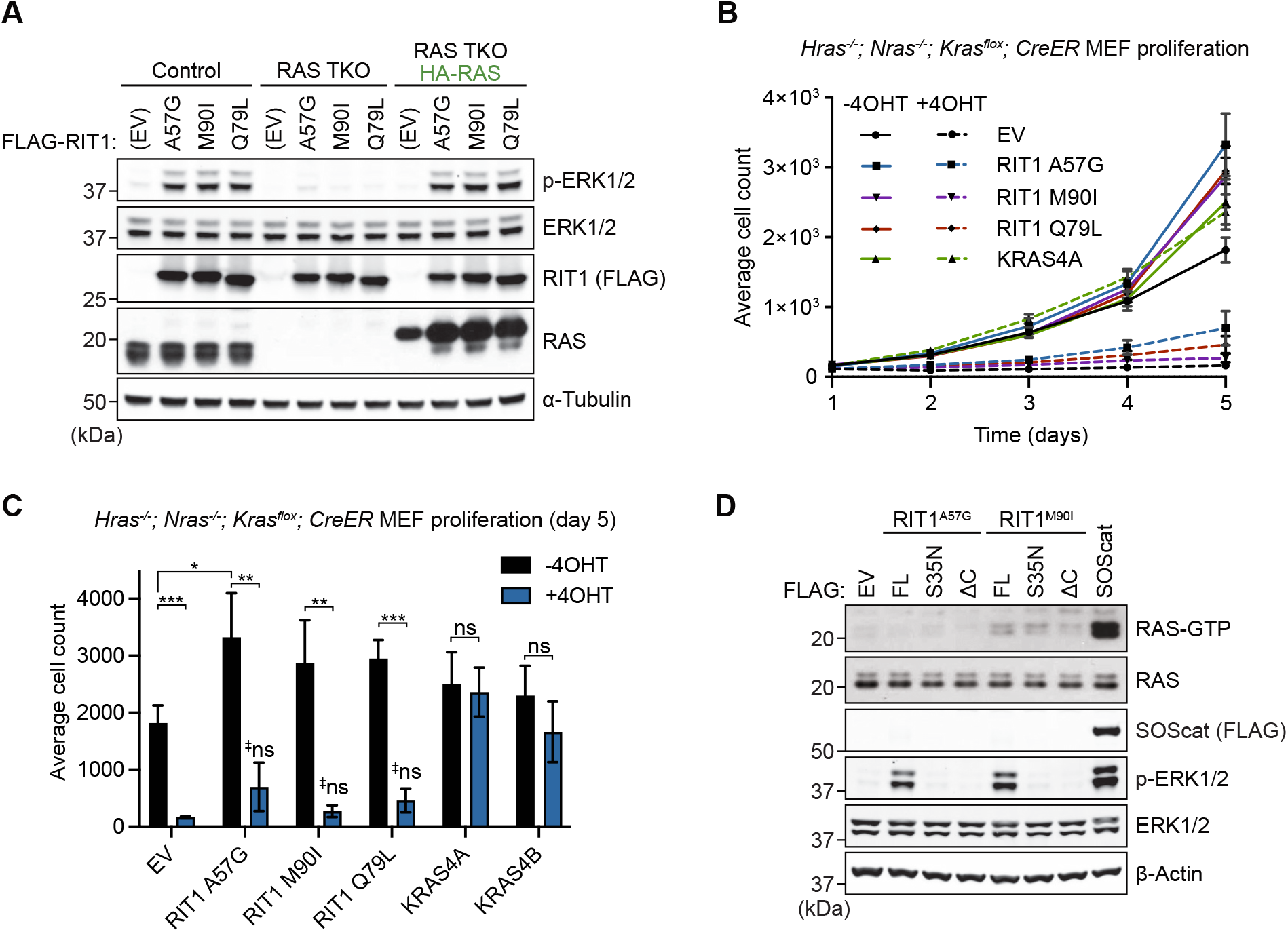
Pathogenic RIT1 relies on RAS to potentiate MAPK signaling. (**A**) Immunoblot analysis of indicated proteins from Rasless (HRAS/NRAS/KRAS TKO) or control HEK293 cells transiently transfected with indicated FLAG-tagged RIT1 constructs or an empty vector (EV) control and serum-starved for 16 h. Rasless cells were rescued with ectopic expression of HA-tagged HRAS, NRAS, KRAS4A, and KRAS4B (1:1:1:1 DNA ratio). One of two independent experiments is shown. (**B**) Proliferation curves of control (−4OHT) and Rasless (+4OHT) MEFs stably expressing indicated constructs. Data points indicate the mean ± SEM of three biological replicates. (**C**) Relative cell growth of control (−4OHT) and Rasless (+4OHT) MEFs stably expressing indicated constructs at Day 5 of growth assay as in C. Data points indicate mean ± SD, *n* = 3, Two-sided Student’s t-test, *p ≤ 0.05, **p ≤ 0.01, ***p ≤ 0.001; ^‡^ns (p>0.05, vs EV+4OHT). (**D**) Immunoblot analysis of indicated proteins from HEK293 cells transiently transfected with indicated FLAG-tagged constructs and serum-starved for 16 h. GTP-bound RAS was precipitated with immobilized RAF1-RBD. SOScat (SOS1 a.a. 564–1049). One of two independent experiments is shown.

As an additional method to assess Ras dependency, we used MEFs that can be rendered “Rasless” upon treatment with 4-hydroxytamoxifen (4OHT) (*35*). Upon genetic deletion of all three Ras genes, pathogenic RIT1 expression failed to restore MAPK pathway activation in response to FBS stimulation compared to ectopically expressed WT KRAS4A or KRAS4B (**fig. S6C**). Since Rasless MEF proliferation is MAPK-dependent (*35*), we assessed cell growth as an additional readout of MAPK activity (**Fig. 7B**). Ectopic expression of mutant RIT1 failed to rescue Rasless cell growth; however, we note that a trend toward a partial rescue, most noticeable with RIT1^A57G^, was observed (**Fig. 7C**). Moreover, mutant RIT1 expression enhanced the proliferation rate of control MEFs, consistent with RIT1-mediated MAPK pathway activation observed here and in other cell models with endogenous Ras expression (*6, 11, 13, 36*). These data suggest that although mutant RIT1 is capable of direct RAF binding, its ability to activate the MAPK pathway is potentiated by the presence of classical Ras proteins. To rule out indirect MAPK activation upstream of Ras (e.g., through the regulation of positive RAS regulators, such as SHP2 or SOS1/2), we measured Ras-GTP levels from cells expressing pathogenic RIT1 and observed no increase in GTP-loaded Ras that correlated with MAPK pathway activation (**Fig. 7D**).

### MEK inhibition attenuates pathological RIT1 MAPK activation and hypertrophy in cardiac tissues

Mutations in pathway components upstream (SHP2 and SOS1) and downstream (RAF1, SHOC2) of Ras that promote MAPK activation define the NS pathogenic landscape (*12*). Intriguingly, the development of NS-associated congenital heart defects, a primary cause of morbidity and mortality for these individuals, varies depending on the genetic driver. This is particularly notable in certain NS genotypes that are more likely associated with HCM, such as RAF1 and RIT1 NS (*9, 12*). The high incidence of HCM (50-70%) and related heart defects (pulmonary stenosis and atrial septal defects) (*9*) in RIT1 NS patients prompted us to establish an in vitro model system to investigate the impact of mutant RIT1 expression in cardiac cells. To this end, we isolated neonatal cardiomyocytes from mice harboring an engineered *Rit1* locus with Cre recombinase-inducible expression of the pathogenic variant *Rit1*^*M90I*^ (*13*). Upon isolation, cardiomyocytes were treated with adenoviruses encoding for Cre recombinase to induce expression of the *Rit1*^*M90I*^ variant. RNA sequencing analysis on day 6 post adenoviral delivery revealed that RIT1^M90I^ expression had elicited broad alterations in the transcriptomic landscape of cardiac myocytes (**Fig. 8A**). These changes included the upregulation of several well-established MAPK target genes (*Ccnd1, Etv4, Egr2, Dusp2*, and *Ereg*), confirming that pathogenic RIT1 regulates MAPK signaling in this cell type (**fig. S7A**)(*37*). In addition, GO and KEGG analyses revealed an enrichment of genes critical for proper cardiac function (**Fig. 8B,C**) and whose dysregulation may contribute to the cardiomyopathy-like phenotype exhibited by RIT1 NS murine models (*13, 14*). Further, these data suggest that upregulation of MAPK signaling by mutant RIT1 may drive the dysregulation of cardiomyopathy-associated genes, such as *Mybpc3* (Myosin-binding protein C), a causal gene representing approximately 20% of HCM patients (*38*), and *Actc1* (cardiac α-actin), among others (**fig. S7B**) (*38, 39*).

**Fig. 8.**
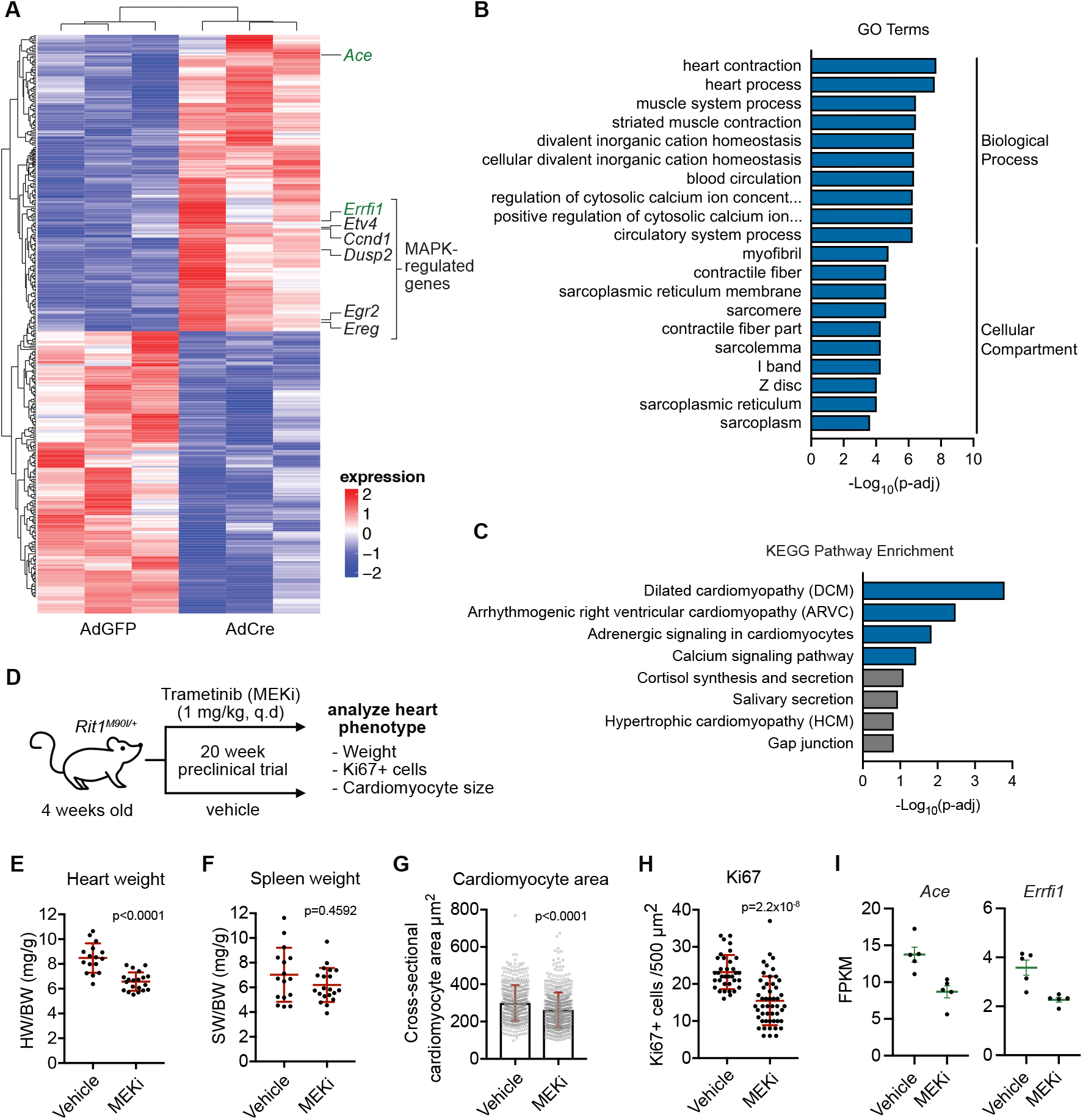
MAPK inhibition alleviates RIT1-dependent cardiac hypertrophy. (**A**) Heatmap of top differentially expressed genes in primary cardiomyocytes from *Rit1*^*LoxP-M90I*^ neonates treated with adenovirus encoding Cre recombinase (AdCre) or GFP (AdGFP). (**B, C**) Gene ontology (GO) and KEGG enrichment analysis of differential gene expression elicited by RIT1^M90I^ expression in primary cardiomyocytes (AdCre vs. AdGFP). (**D**) Schema of 20-week Trametinib (MEKi) preclinical trial with *Rit1*^*M90I/+*^ mice. (**E, F**) Comparison of normalized heart (E) or spleen (F) weight between MEKi (*n* = 16) and vehicle control (*n* = 20) group. Statistical significance was assessed by a two-tailed Mann-Whitney test. Error bars indicate mean ± SD. (**G, H**) Quantification of myocyte area (G) and Ki67+ cells (H) from heart cross sections by immunofluorescence and immunohistochemistry, respectively. Statistical significance was assessed by a two-tailed Mann-Whitney test. Error bars indicate mean ± SD. (**I**) Normalized mRNA transcript levels (FPKM) of indicated genes in hearts (*n* = 5) isolated from vehicle control or MEKi-treated *Rit1*^*M90I/+*^ mice at 20-week endpoint. Error bars indicate mean ± SEM.

Given the data above, we leveraged our RIT1 NS mouse model to evaluate whether pharmacological inhibition of the MAPK pathway may ameliorate RIT1^M90I^-driven cardiac tissue hypertrophy (*13*). To this end, we treated a cohort of 4-week-old mice harboring a germline *Rit1*^*M90I*^ variant with the allosteric MEK1/2 inhibitor trametinib (MEKi) or vehicle control (**Fig. 8D**). After 20 weeks of daily treatment, we observed a significant decrease in heart weight of MEKi-treated mice (**Fig. 8E**), but no difference in spleen weight (**Fig. 8F**), suggesting that MEK inhibition may reduce aberrant cardiac tissue growth associated with mutant RIT1 expression. Indeed, when the size (cross-sectional area) and proliferative state (Ki67 staining) of myocytes from MEKi and control hearts were compared, MEKi-treated hearts exhibited a marked reduction in both parameters, indicating reduced cell growth (**Fig. 8G, H**). Transcriptomic profiling by RNA-seq confirmed that systemic MEKi treatment effectively inhibited MAPK signaling and suggests that the observed reduction in cardiac cell growth was a direct consequence of MEK inhibition within *Rit1*^*M90I/+*^ hearts (**fig. S7C**). Together, these data suggest that pharmacological inhibition of aberrant RIT1-mediated MAPK signaling may represent a viable therapeutic strategy to ameliorate the cardiac defects presented by RIT1 NS patients. Further analysis of our differential gene expression datasets identified two genes, *Ace* (Angiotensin 1 converting enzyme) and *Errfi1* (ERBB Receptor Feedback Inhibitor 1), which were transcriptionally upregulated upon RIT1^M90I^ expression in primary cardiomyocytes, see Fig. 6A, and downregulated in *Rit1*^*M90I/+*^ hearts following MEKi treatment (**Fig. 8I**). These patterns of expression mark *Ace* and *Errfi1* as potential biomarkers for RIT1 NS individuals with associated HCM.

## DISCUSSION

Activation of the MAPK pathway occurs at the inner leaflet of the PM wherein lipid-anchored Ras GTPases recruit RAF kinases and facilitate a multi-step activation process resulting in active RAF dimers (*30*). Since the discovery of RIT1, and its paralog RIT2, the absence of HVR prenylation motifs prompted early speculation into their unique HVR-dependent PM association (*40*). Here, we show that RIT1, like the classical Ras GTPases, requires membrane binding for pathological MAPK activation. An extended polybasic HVR, containing three PBRs, mediates electrostatic interaction with negatively charged phospholipids, a property akin to the polybasic KRAS4B HVR; however, the absence of a RIT1 HVR lipid anchor may allow for transient and dynamic association with the PM. We have shown that RIT1 diffuses between PM and cytoplasm during mitosis to interact with spindle assembly checkpoint proteins MAD2 and p31^comet^, a process that is regulated by CDK1-mediated HVR phosphorylation (*31*). Furthermore, appending a C-terminal prenylation motif prevents dissociation from the PM and blocks RIT1 mitotic regulation. Intriguingly, we and others (*23*) have identified non-charged residues (Trp^204^, Leu^207^, Phe^211^) interspersed between the PBRs critical for membrane association. Molecular dynamics simulations have revealed that the hydrophobic side chains of these residues burry deeply into the lipid bilayer (*22*); however, further investigation is needed to determine whether these residues help coordinate the association of PBRs with phospholipid head groups. We speculate that the uniquely electrostatic association with the PM may enable RIT1 to sense the composition of inner leaflet lipids.

To best interrogate the contribution of membrane association with RAF binding, we developed a BRET assay to quantitate RIT1-RAF association in the context of a native PM environment. We found that perturbations of the RIT1 HVR that abrogated membrane association strongly correlated with decreased RAF binding. Although anticipated, these data exemplify the critical nature of the RIT1 HVR in mediating RIT1’s diverse functions. In contrast with previous reports demonstrating that RIT1 associates preferentially with BRAF in neuronal cells (*17*), we found that RIT1 binds most strongly with RAF1 in vitro and in intact cells, suggesting that selectivity towards RAF isoforms may be context-dependent. Interestingly, preferential RAF1 binding is a property also shared by the classical Ras family members in cells (*24*).

Accumulation of RIT1 through the loss of LZTR1-mediated proteasomal degradation increases MAPK signaling, a defining pathognomonic feature of RASopathies (*13, 15, 41*). However, as with other NS germline mutants, pathologic RIT1 signaling is mild and thus compatible with embryonic development (*1*). Despite reporting by independent groups of a direct, albeit weak, interaction between RIT1 and RAF kinases (*13, 17, 42*), evidence of RAF activation was limited. The data presented here identify RAF kinases, namely RAF1, as the primary effectors through which mutant RIT1 proteins activate MAPK signaling.

Nonetheless, it remains possible that RIT1 activates ERK1/2 through alternate MAP3 kinases (*36*), and merits further investigation to determine whether different cell contexts determine effector selectivity. Intriguingly, we have found that activation of MAPK still requires classical RAS proteins, consistent with the fact that deletion of these proteins in mouse cells results in a complete growth arrest (*35*). Further, mutant RIT1 expression did not influence the proportion of GTP-loaded Ras, suggesting that RIT1 promotes MAPK pathway activation downstream of Ras. We posit that despite the low RIT1-RAF affinity, the overabundance of mutant RIT1 protein may facilitate Ras-RAF activation by increasing the local concentration of RAF at the PM, thereby “priming” Ras-RAF activation in response to upstream RTK signaling (**Fig. 9**). However, further investigation is needed to shed light on the exact mechanism. Nonetheless, our findings suggest that RIT1-driven disease may be treated not only with MAPK pathway inhibitors but also with inhibitors that limit upstream Ras activation (e.g., SOS1 and SHP2 inhibitors).

**Fig. 9.**
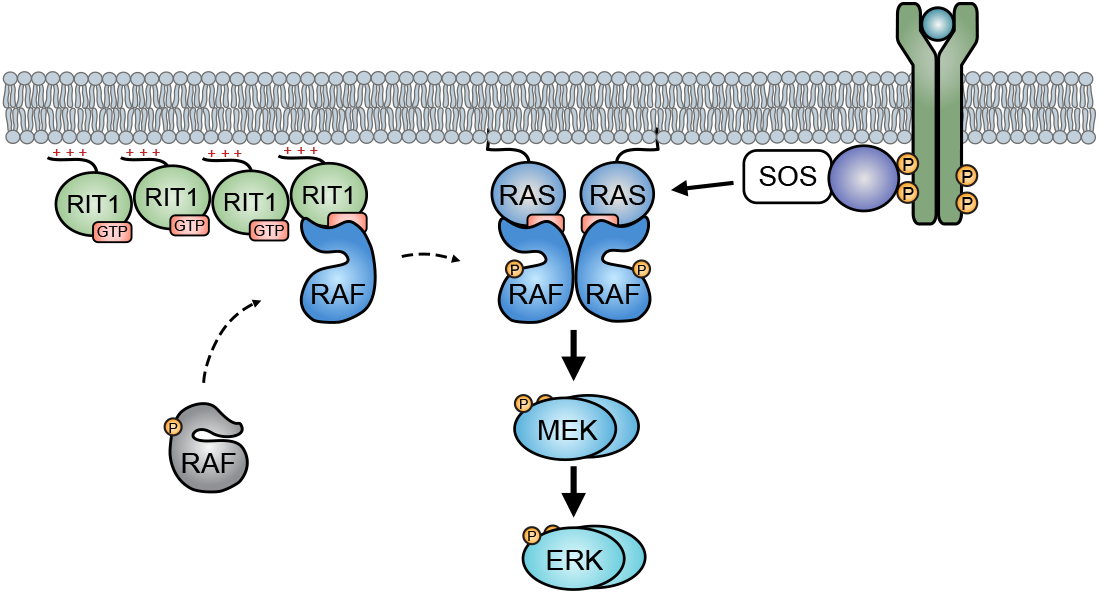
Schematic model of Ras-dependent RIT1-mediated MAPK hyperactivation. Accumulation of proteolysis-deficient, pathogenic RIT1 protein recruits RAF kinases to the PM thereby “priming” complete RAF activation by PM resident GTP-loaded Ras.

Approximately one-fifth of individuals with RIT1 NS harbor an A57G allele, making it the most common RIT1 variant in this condition (*8*). RIT1^A57G^ is of particular interest, biochemically, due to its hypermorphic enhanced binding to RAF kinases shown here while exhibiting comparable rates of intrinsic GTP hydrolysis, nucleotide exchange, and cellular fraction bound to GTP as other gain-of-function alleles (*13, 42*). Intriguingly, RIT1^A57G^ activates MAPK signaling to a similar extent as other pathogenic variants despite an increased affinity to RAF (*13, 33*). Thus, RIT1 activation of RAF is not solely correlated to the strength of their interaction, suggesting this is not the limiting factor associated with the modest degree of MAPK pathway activation exhibited by pathogenic RIT1 variants. The cysteine-rich domain (CRD) of RAF1 makes critical contacts with the inter-switch region of KRAS that are essential for RAF activation (*25*). We speculate that the inter-switch region of RIT1, which shares low homology with the Ras inter-switch region, likely fails to engage productively with the CRD of RAF.

Compared to patients with other genetic variants, RIT1 NS individuals exhibit an elevated frequency of HCM (*9, 12*). In a prior study, we described a RIT1^M90I^ mouse model that recapitulated clinical manifestations of NS disease, including cardiac hypertrophy (*13*). Here, we demonstrate that treatment of RIT1^M90I^ mice with the FDA-approved MEK1/2 inhibitor trametinib (GSK1120212) ameliorated cardiac tissue overgrowth, suggesting that targeting the MAPK pathway may be an effective therapeutic strategy in NS patients with mutant RIT1. Indeed, off-label trametinib treatment was recently shown to reverse myocardial hypertrophy in three children with RIT1 NS (*43, 44*). However, trametinib has been reported to induce significant levels of toxicity in other disease contexts (*45, 46*), highlighting the need for further pre-clinical work to address optimal dosing and treatment windows, response to different MEK inhibitors, and efficacy of upstream Ras inhibition to alleviate the cardiac and extracardiac RIT1 phenotype. Altogether, these findings aid our mechanistic understanding of RIT1 disease and support the evaluation of broader therapeutic strategies.

Lastly, we must consider the implications of a Ras-dependent RIT1 MAPK activation in the context of RIT1-driven tumors. Cells expressing RIT1^M90I^, but not those expressing KRAS^G12V^, depend on RTK and adaptor proteins upstream of Ras, including EGFR, GRB2, SHP2, and SOS1 for growth (*47*). Conversely, loss of NF1 and SPRED1, two negative regulators of Ras activity, promote mutant RIT1 cell growth, consistent with our model in which RIT1 relies on active Ras to promote MAPK signaling. Intriguingly, in the same system, Vichas et al. (2021), show that loss of LZTR1 similarly promotes the growth of cells expressing ectopic RIT1^M90I^, a variant that escapes LZTR1-mediated degradation, suggesting that stabilization of endogenous RIT1 may provide an additional growth advantage (*13*). In certain cell contexts, LZTR1 promotes the degradation of other Ras family GTPases, most notably MRAS (*13, 48*), and its loss may enhance MRAS-mediated RAF activation in synergy with mutant RIT1 expression (*49*). In support of this hypothesis, mutant RIT1 cells exhibited a strong growth dependency on SHOC2, a scaffolding protein that associates with MRAS to promote Ras-dependent activation of RAF (*50*). Additionally, RIT1 moonlights as a mitotic checkpoint regulator (*31, 47*), among other functions (*8, 33*), imparting mutant RIT1 cancer cells with unique therapeutic vulnerabilities (*47*). While our work sheds some light on RIT1’s dependency on RTK-MAPK components, further studies are needed to define the contribution of RIT1’s various functions to its oncogenic potential.

## MATERIALS AND METHODS

### Protein production

#### Cloning

DNA constructs for the expression of KRAS4B (1-169) (Addgene #159539) and RAF1 (52-131) were previously described (*50*). Gateway Entry clones for BRAF (151-277), RIT1 (17-197), RIT1 A57G (17-197), and RIT1 (17-219) were generated by standard cloning methods and incorporate an upstream tobacco etch virus (TEV) protease cleavage site (ENLYFQG) followed by the appropriate coding sequences. Sequence-validated Entry clones were sub-cloned into pDest-566, a Gateway Destination vector containing a His6 and maltose-binding protein (MBP) tag to produce the final *E. coli* expression clones (*51*). All expression constructs were in the form containing an N-terminal His6 and maltose-binding protein (MBP) tag (Addgene #11517).

#### Protein expression

RAF1-RBD (52-131) was expressed using the auto-induction media protocol and BRAF (151-277) and RIT1 proteins were expressed using the Dynamite media protocol (*51*). For ^15^N isotopic incorporation into RAF1 (52-131), KRAS4B (1-169), RIT1 (17-197), RIT1 (17-219), and RIT1 (17-197) A57G, seed cultures were inoculated from glycerol stocks of the transformed strains into 300 mL of Studier’s MDAG135 medium (*52*) (25 mM Na_2_HPO_4_, 25 mM KH_2_PO_4_, 50 mM NH_4_Cl, 5 mM Na_2_SO_4_, 2 mM MgSO_4_, 50 µM FeCl_3_, 20 µM CaCl_2_, 10 µM MnCl_2_-4H_2_O, 10 µM ZnSO_4_-7H_2_O, 2 µM CoCl_2_-6H_2_O,

2 µM CuCl_2_-2H_2_O, 2 µM NiCl_2_-6H_2_O, 2 µM Na_2_MoO_4_-2H_2_O, 2 µM Na_2_SeO_3_-5H_2_O, 2 µM H_3_BO_3_), 19.4 mM glucose, 7.5 mM aspartate, and 200 µg/ml each of 18 amino acids (E, D, K, R, H, A, P, G, T, S, Q, N, V, L, I, F, W, M) in a 2 L baffled shake flask for 16 hours at 37°C until late-log phase growth. In the interim, 15 L of T-20052 (*53*) medium was prepared in a 20-liter Bioflow IV bioreactor (Eppendorf/NBS). The seed culture was collected and centrifuged at 3,000 x *g* for 10 minutes at 25¼;C. The pellet was re-suspended with 100 mL of the sterilized T-20052 medium from the bioreactor and then returned to the bioreactor as inoculum. The culture was grown at 37°C with an airflow of 15.0 LPM and agitation of 350 RPM. Approximately five hr (mid-log phase) after inoculation, the culture was shifted to 20°C overnight. Cells were collected by centrifugation (5000 x *g* for 10 min at 4°C) and the cell pellet was stored at -80°C.

#### Protein purification

Proteins were essentially purified as previously described for proteins in the His6-MBP-tev-POI (protein of interest) expression format (*54*). Note that 5 mM MgCl_2_ was used in buffers used to purify KRAS4B (1-169). Essentially, the His6-MBP-POI was purified by immobilized metal-ion affinity chromatography (IMAC) from the lysate, the His6-MBP tag was removed by His6-TEV protease digestion, the POI was isolated from the TEV digest by another round of IMAC (POI in flow-through, wash, or low imidazole elutions), the pooled protein was buffered exchanged via preparative SEC, concentrated and frozen in liquid nitrogen in aliquots, and stored at -80°C. Purifications of RIT1 required some alterations to this basic protocol. Specifically, protein concentration was kept below 4 mg/ml throughout, all ^15^N preparations of RIT1 (17-197) and preparations of RIT1 (17-219) were with 300 mM NaCl and 10% (w/v) glycerol in all buffers. Final buffers were 20 mM HEPES, pH 7.3, 150 mM NaCl, 1 mM TCEP for BRAF (151-277), 10 mM Tris-HCl, pH 7.5, 50 mM NaCl, 2 mM MgCl_2_ for RAF1 (52-131), 20 mM HEPES, pH 7.3, 150 mM NaCl, 2 mM MgCl_2_, 1 mM TCEP for RIT1 (17-197) and KRAS4B (1-169) with 5 mM MgCl_2_ added for KRAS concentrations greater than 10 mg/ml, and 20 mM HEPES, pH 7.3, 300 mM NaCl, 2 mM MgCl_2_, 1 mM TCEP, 10% glycerol for RIT1 (17-219) and ^15^N preparations of RIT1 (17-197).

### Liposome SPR

2.5 mM Liposomes were prepared from various amounts of 1-palmitoyl-2-oleoyl-glycero-3-phosphocholine (POPC) and 1-palmitoyl-2-oleoyl-sn-glycero-3-phospho-L-serine (POPS). Lipid mixtures were lyophilized at -80°C for approximately three hours. Lipids were reconstituted in 1 ml 20 mM HEPES pH 7.4, 150 mM NaCl buffer and sonicated at 37°C for 5 minutes. Mixtures underwent five freeze/thaw cycles, followed by another brief sonication until clear.

SPR binding experiments were run on a Biacore T200. Liposomes composed of POPC or POPS were captured on flow cells 2-4 of an L1 Chip, flow cell 1 was unmodified and served as a reference. RIT1 was diluted in SPR running buffer (20 mM HEPES pH 7.4, 150 mM NaCl), concentration range, 5 – 0.156 µM (1:2-fold dilutions) and injected on the chip surface at 30 µl/min. Sensorgrams were normalized by the capture level of liposomes. Partition coefficients were calculated as described previously (*55*).

### GNP exchange

RIT1 (2 mg) was diluted in Alkaline Phosphatase buffer (40 mM Tris pH 7.4, 200mM Ammonium Sulphate, 1mM ZnCl_2_, 10% glycerol). 10 mM GppNHp and 5 units of alkaline phosphatase beads were added to RIT1, and the mixture was incubated for one hour at 4°C with constant rotation. Beads were pelleted out, and 30 mM MgCl_2_ was added to the mixture, followed by another brief incubation at 4°C. Protein was desalted on a HiPrep 26/10 desalting column into buffer (20 mM HEPES pH 7.4, 150 mM NaCl, 5 mM MgCl_2,_ 1 mM TCEP). The efficiency of exchange was measured by HPLC analysis, as previously published (*53*).

### Isothermal titration calorimetry

ITC experiments were run on a MicroCal PEAQ-ITC instrument. The proteins were diluted to 100 µM (RIT1) and 300 µM (RAF) in 20 mM HEPES, 150 mM NaCl, pH 7.4, 5 mM MgCl_2_, 1 mM TCEP buffer. Approximately 200 µl RIT1 was loaded into the cell of the instrument, and 80 µl RAF was loaded into the syringe. 4.4 µl of RAF1 was titrated onto RIT1 every 2 minutes, for a total of 19 injections. After all injections were complete, the data was analyzed using the MicroCal PEAQ-ITC analysis software to calculate the *K*_*D*_.

### NMR

RAF1-RBD (52-131), RIT1(17-197) WT/A57G, and WT KRAS4B (2-169) and their complexes were characterized by solution NMR spectroscopy (see Appendix Fig S1-S7). For all these proteins, published backbone assignments were used initially with in-home NMR assignments whenever required (BMRB IDs: RAF1 (17382), RIT1 (26787), and KRAS (28021)). All ^15^N labeled proteins used in this study had concentrations between 100 µM to 150 µM. Protein-protein complexes were pre-formed at a 1:3 ratio (3-fold excess of unlabelled partner) and the saturating complex was confirmed by observing the signal of the key reporting residues. 2D ^1^H-^15^N HSQC spectra were recorded at 25 °C on a Bruker 700MHz spectrometer equipped with proton-cooled cryogenic ^1^H/^13^C/^15^N triple resonance probes. The sample temperature was calibrated with a 100% methanol sample before the experiments. All ^1^H – ^15^N HSQC spectra were collected with 1K and 128 complex points in F2 (^1^H) and F1(^15^N) respectively, with 16 scans. All experiments used a ^1^H spectra width of 9090 Hz and a ^15^N spectral width of 1987 Hz with the proton and ^15^N carriers set to 4.7 ppm and 120 ppm. Data were processed by NMRPipe (*56*), analyzed by NMRFAM-SPARKY (*57*), GraphPad Prism was used to plot CSP plots and PyMOL was used to map CSP to X-ray structures. Chemical shift perturbations (CSP) were calculated by using the equations Sqrt{(dH^^2^+(dN/10)^^2^)/2}, where dH and dN are the proton and nitrogen chemical shift differences between the complexed and non-complexed proteins.

### Cell lines and culture conditions

HeLa and HEK293T cells were obtained from the American Type Culture Collection (ATCC). FlpIn HEK293 cells were obtained from Thermo Fisher. *Hras*^*-/-*^; *Nras*^*-/-*^; *Kras*^*flox*^; *CreER* mouse embryonic fibroblasts (MEFs) were a kind gift from Nikki Fer (Frederick National Laboratory). Cells were cultured in Dulbecco’s modified Eagle’s medium (DMEM) supplemented with 10% Fetal Bovine Serum (FBS). Cells were grown in a humidified incubator with 5% CO2 at 37°C. Validation procedures are as described by the manufacturer. Cell lines were regularly tested and verified to be mycoplasma negative using MycoAlert PLUS Mycoplasma Detection Kit (Lonza).

For the generation of Rasless 293 cells, crRNA targeting HRAS (Integrated DNA Technologies (IDT)) and ATTO 550 labeled tracrRNA (IDT) were combined to a final concentration of 1 µM and annealed by cooling from 95°C to room temperature. 12 pmol crRNA:tracrRNA duplex was combined with 12 pmol recombinant HiFi Cas9 (IDT) and reverse transfected into HEK293 Flp-In cells using Lipofectamine CRISPRMAX reagent (Thermo Fisher Scientific) following IDT protocols. 24 hours after transfection ATTO 550 positive cells were sorted using a Sony SH800 Cell Sorter and allowed to recover for four days. Following the same procedure, cells were then sequentially transfected with NRAS and KRAS crRNA:tracrRNA-Cas9 ribonucleoprotein complexes. For the final cell sorting step following KRAS crRNA:tracrRNA-Cas9 transfection, single ATTO 550 positive cells were sorted into 96-well plates. Clonal cell lines were then expanded and screened for HRAS, KRAS and NRAS knock-out by western blotting using antibodies specific for the individual RAS isoforms and two pan-H/K/NRAS antibodies. Five confirmed triple knock-out clones were then pooled to mitigate off-target guide RNA effects and clonal heterogeneity, and validated functionally through the absence of MAPK signaling. To generate HEK293 Flp-In control sgRNA cells, the same procedures were performed in parallel targeting safe loci in AAVS1, chromosome 3 and chromosome 15 (*58*). Single-cell clones were expanded, and normal activity and expression of RAS signaling pathway components were confirmed by western blotting. 12 clones were then pooled to give a polyclonal control sgRNA cell line. See Supplementary Table S2 for sgRNA targeting sequences.

For the generation of Rasless MEFs with stable expression of ectopic RIT1 or Ras, lentivirus was produced by co-transfection of HEK293T cells with a lentiviral vector and the packaging plasmids psPAX2 (Addgene, plasmid #12260) and pMD.2G (Addgene, plasmid #12259) at a ratio of 1.25:1.0:0.25. The supernatant was collected 72 hours post-transfection and filtered through a 0.45 µm filter. *Hras*^*-/-*^; *Nras*^*-/-*^; *Kras*^*flox*^; *CreER* MEFs were transduced with lentiviral-containing supernatant supplemented with 0.8 µg/ml polybrene (Sigma-Aldrich). Stably transduced cells were selected with 1.5 µg/ml Puromycin (Sigma-Aldrich). To remove the floxed *Kras* allele, stable cells were treated with 1µM 4-hydroxytamoxifen (4OHT, Sigma-Aldrich). Assays with Rasless MEFs were conducted 10-11 days post 4OHT treatment and loss of KRAS was verified by immunoblot.

### Mammalian expression constructs

All RIT1 cDNA mutants were generated using standard PCR-based site-directed mutagenesis in the pDONR223-RIT1 template and were previously described (*31*). KRAS4A and KRAS4B entry clones (Addgene plasmids 83166 and 83129) and ARAF, BRAF, and RAF1 (plasmids 70293, 70299, and 70497) were a gift from Dominic Esposito (Frederick National Lab). SOScat (SOS1 residues 564–1049) was previously described (*15*). For N-terminal GST-tagged proteins, entry clones were gateway cloned into pDEST27 destination vector (Invitrogen). For N-terminal mCherry-, EGFP- or FLAG-tagged constructs, entry clones were cloned by multisite gateway cloning into pDEST302 or pDEST663 (a gift from Dominic Esposito, Frederick National Lab) and with expression controlled by an EF1a promoter. HA-tagged constructs were gateway cloned into a pcDNA-HA destination vector. Empty vector (EV) plasmid controls were generated using a gateway recombination cassette containing a stop codon followed by an untranslated stuffer sequence.

### RNA interference

To knockdown RAS, individual pools of short interfering RNAs (siRNAs) against mouse *Hras, Nras*, and *Kras* (SMARTpool: ON-TARGETplus, Horizon) were pooled in equal amounts and transfected into cells using Lipofectamine RNAiMax Transfection Reagent (Life Technologies) according to manufacturer’s instructions. Similarly, knockdown of RAF kinases was achieved using siRNA targeting human *RAF1, ARAF*, and *BRAF* (SMARTpool: ON-TARGETplus, Horizon). Non-targeting control siRNA was purchased from Horizon (D-001810-10-05).

### Primary cardiomyocyte isolation

Preparation of mouse neonatal cardiac myocytes was conducted as previously described (*59*) with some modifications. Briefly, hearts were extracted from 1–3-day old neonates, placed in ice-cold HBSS, cleared of blood clots and aortic tissue, then placed in fresh ice-cold HBSS and cut into small pieces to be digested in prewarmed HBSS with 1 mg/ml Collagenase Type 2 (Life Technologies) at 37 °C for 5 min with gentle agitation. Heart pieces were then allowed to sediment for 1 minute, and the supernatant was removed and discarded. The following digestion process was repeated six times: heart pieces from before were resuspended in fresh HBSS with 1 mg/ml Collagenase and incubated at 37 °C for 5 min with gentle agitation. Heart pieces were then allowed to sediment for 1 min and the supernatant containing suspended cardiac myocytes was transferred to a new tube containing 1/10 volume of cardiomyocyte medium (DMEM supplemented with 10% FBS, 10% Nu serum (Corning), 1x Penicillin-Streptomycin (Gibco), 1x Glutamax (Gibco), 1x ITS liquid medium supplement (Sigma), and 10 mM HEPES (Gibco)) and centrifuged to pellet cells (500g for 5 min). Cell pellets were resuspended in 1 ml cardiomyocyte medium and pooled after the sixth collection, passed through a cell strainer multiple times and incubated at 37 °C for 2 hours in a plastic 10cm dish. Unadhered cells in cell suspension were then pelleted by centrifugation and resuspended in 1 ml cardiomyocyte medium per heart and seeded on a laminin-coated 12-well plate at a density of ∼1 heart per well. Cells were exchanged into fresh media daily and infected with 2 × 10^8 adenoviral particles encoding GFP or GFP-Cre (ViraQuest), per well, on day 3.

### RT-qPCR

Total RNA from cardiomyocytes was obtained on day 6 post adenovirus infection using the RNeasy kit (Qiagen) according to the manufacturer’s instructions. cDNA was obtained by reverse transcription (RT) of 1 ug RNA using qScript XLT cDNA SuperMix (QuantaBio; 95161). 10 ng of cDNA was diluted in nuclease-free water and ran in technical triplicates using PowerUp SYBR Green Master Mix (Applied Biosystems) on a QuantStudio 5 (Thermo Fisher Scientific). *Tbp* (TATA-box binding protein) was used as an endogenous control. See Supplementary Table S3 for primer sequences.

### Live-cell imaging

HeLa cells were seeded onto 12-well #1.5 glass bottom plates (Cellvis) and transiently transfected with Fugene 6 (Promega), following the manufacturer’s instructions. Before imaging, cells were exchanged into imaging media: FluoroBrite DMEM (Thermo Fisher Scientific) supplemented with 10% FBS and 4 mM GlutaMAX (Thermo Fisher Scientific). Images were acquired as a series of 0.6 µm z-stacks with a Plan Apo 40x/0.95 Corr (DIC N2 / 40X I) 227.5 nm/pixel objective (Nikon) on a Nikon Ti-E inverted CSU-22 spinning disk confocal microscope equipped with an incubation chamber (Okolab), providing a humidified atmosphere at 37 °C with 5% CO2. Images were processed using Fiji (*60*).

### NanoBRET RIT1/RAF Interaction Assay

HEK293T cells were seeded in a 12-well plate at 1.25 x105 cells/well density. After 24 hours, cells were transfected with mVenus-RIT1 and RAF-nanoLuc using Fugene 6. The concentration of donor (nanoLuc) was kept constant, and the concentration of acceptor (mVenus) was diluted 2-fold (concentrations range 0ng-1000 ng). An empty vector plasmid was transfected into cells to normalize the DNA amount in each well. 48 hours post-transfection, cells were trypsinized and recovered in DMEM cell media containing 10% fetal bovine serum. Tubes were spun down at 1500 rpm for 3 minutes and the pellet was resuspended in Dulbeccos’s PBS (PBS) + 0.5% FBS. A cell suspension (20,000 cells) was added in triplicate to both a white 384-well PE Optiplate (for BRET reading), and a black 384well plate (for mVenus reading). 20 PBS + 0.5% FBS was added to each well of cells in the black plate to bring the final volume to 40 µl. 20 µl of 30 mM nanoBRET nano-Glo substrate (Promega) was added to all required wells of the 384-well PE Optiplate. Plates were read on a PerkinElmer Envision plate reader. The white plate was monitored at 535 nm (BRET signal) and 470 nm (background nanoLuc). mVenus was monitored in the black plate at 530 nm emission with an excitation at 500 nm. The BRET value at each point was measured by dividing the BRET signal by the background nanoLuc signal. Acceptor/donor ratios were normalized against control with equal amounts of mVenus-RIT1(1-219) and RAF-nanoLuc transfected. Data were analyzed using a non-linear regression fit with GraphPad Prism software to obtain the BRET_50_ values.

### Pulldowns and Immunoblotting

For GST pulldown of proteins from cell lysates, 3 × 10^6^ HEK293T cells were transfected with 4 µg total DNA of indicated plasmids. 24 hours after transfection, cells were rinsed with ice-cold PBS and lysed with 1 ml of lysis buffer (50 mM Tris-HCl (pH 7.5), 150 mM NaCl, 1% IGEPAL CA-630, 10% glycerol) supplemented with protease and phosphatase inhibitor cocktails (Sigma-Aldrich). Lysates were cleared by centrifugation and incubated with 20 µl of Glutathione Sepharose 4B beads for 4 h at 4°C with end-over-end rotation. Beads were rinsed three times with Lysis buffer and resuspended in LDS sample buffer.

Whole-cell lysates for immunoblot analysis were prepared using RIPA Buffer (50 mM Tris-HCl (pH 8.0), 150 mM NaCl, 0.5% sodium deoxycholate, 0.1% SDS, 1% IGEPAL CA-630) supplemented with protease and phosphatase inhibitor cocktails (Sigma-Aldrich). 15-30 µg of total protein was loaded per well of precast NuPAGE gels (Life Technologies).

For immunoblot detection, samples were separated by SDS-PAGE and transferred onto nitrocellulose membranes. Membranes were blocked using 5% skimmed milk in TBST buffer for 1 hour and incubated with appropriate primary antibodies overnight. Detection was performed using secondary antibodies conjugated to DyLight680 (611-144-002; 1:10,000) or DyLight800 (610-145-002; 1:10,000) (Rockland), and visualized with a LI-COR Odyssey infrared scanner or using HRP-linked secondary antibodies and developed with Amersham ECL (Cytiva Life Sciences) and X-ray films. Primary antibodies against p-ERK (4370; 1:1000-2000), ERK1/2 (4696 and 4695; 1:1000), p-MEK (9154; 1:1000), MEK1/2 (4694 and 8727; 1:1000), p-EGFR (3777; 1:1000), EGFR (4267; 1:1000), HA (3724, 1:1000), and FLAG (14793; 1:1000) were obtained from Cell Signal Technology. Antibodies against GST (sc-138; 1:1000) and HRAS (sc-520; 1:500) were obtained from Santa Cruz Biotechnology. RIT1 (ab53720; 1:1000), NRAS (ab167136; 1:2000), and pan-RAS (ab108602; 1:1000) antibodies were from Abcam. KRAS (WH0003845M1; 1:500), βActin (A2228; 1:10000), ɑ-Tubulin (T6199; 1:5000), and FLAG (F1804; 1:2000) antibodies were purchased from Sigma-Aldrich.

### Kinase Assay

To measure RAF activity in vitro, FLAG-tagged RAF1 or BRAF was co-expressed with RIT1^A57G^ or empty vector control in HEK293T cells. 24 h after transfection, cells were rinsed with ice-cold PBS and lysed with 1 ml of lysis buffer (50 mM Tris-HCl (pH 7.5), 150 mM NaCl, 1% IGEPAL CA-630, 10% glycerol) supplemented with protease and phosphatase inhibitor cocktails (Sigma-Aldrich). Lysates were cleared by centrifugation and incubated with 10 µl of anti-FLAG M2 agarose beads (EMD Millipore) for 4 h at 4°C with end-over-end rotation. Beads were rinsed three times with lysis buffer and twice with TN buffer (20 mM Tris pH 7.5, 150 mM NaCl, 5% Glycerol). RAF protein was then eluted with 200 μg/mL 3xFLAG peptide (Sigma-Aldrich) in TN buffer, aliquoted, and snap-frozen in liquid nitrogen. Kinase reactions were performed as follows: 100 µl reactions containing 2.5 nM FLAG-RAF1 or 1 nM FLAG-BRAF protein and 1 µM recombinant Avi-MEK1 (K97R), a gift from Dom Esposito (Frederick National Lab), 20 mM Tris pH 7.5, 100 mM NaCl, 1 mM MgCl2, 1 mM DTT and 0.5 mM ATP. 20 µl fractions were removed at indicated time points and the reaction was stopped by the addition of 2x LDS sample buffer.

### Mice

Conditional Rit1^M90I^ mice were previously described (*13*). To generate the experimental cohorts, conditional homozygous male Rit1^M90I^ mice were crossed to homozygous female CMV-Cre deleter transgenic mice. Offspring were weaned at three weeks of age and treatment was started at four weeks of age for a period of 20 weeks. Both male and female littermates were included in this study. Trametinib was purchased from Selleckchem and was diluted in 0.5% carboxymethylcellulose and 0.2% Tween-80 (Sigma). Upon completion of the 20 weeks, mice were given the last dose in the morning and euthanized 2 hours later. Body, heart, and spleen weight was recorded. Both heart and spleen were fixed in phosphate-buffered formalin overnight. This study was performed in accordance with the guidelines in the Guide for the Care and Use of Laboratory Animals of the National Institutes of Health. All the animals were handled according to approved institutional animal care and use committee (IACUC) protocol #AN165444 of the University of California San Francisco.

For the quantification of cardiomyocyte size, transverse cardiac tissue sections were stained with Texas-Red-conjugated wheat germ agglutinin (W21405; 1:200) to label the cell boundaries and counter-stained with DAPI. Five images of cardiac tissue adjacent to the left ventricle were captured at 20X magnification for at least three mice per treatment group. The cross-sectional area of 30 cardiomyocytes per image was measured using Fiji.

## Supplementary Materials

Fig. S1. In vitro RAF kinase assay with RIT1A57G co-expression

Fig. S2. In vitro and in vivo analysis of RIT1-PM lipid associations

Fig. S3. RIT1 BRET assay reveals preferential binding to RAF1 isoform

Fig. S4. NMR spectra of RIT1-RAF1 RBD complexes

Fig. S5. RIT1 C-terminus is essential for PM association but not GTP loading

Fig. S6. RIT1 protein stabilization fails to promote MAPK activation in the absence of RAS

Fig. S7. Analysis of gene expression elicited by RIT1M90I expression or MEK inhibition

Table S1. Broadened residues (blue) and residues above 1.5s reported in CSP plots

Table S2. gRNA targeting sequences

Table S3. qPCR primers

## Acknowledgments

We thank Dom Esposito and Vanessa Wall (Frederick National Lab) for the multisite gateway plasmid toolkit and members of the McCormick lab for their input. We would like to acknowledge Julia Cregger, Matt Drew, Peter Frank, José Sánchez Hernández, Min Hong, Jennifer Mehalko, Zhaojing Meng, Nitya Ramakrishnan, Rosemilia Reyes, Troy Taylor, and Vanessa Wall from the Protein Expression Laboratory at the Frederick National Laboratory for cloning, expression, purification and quality control of all proteins used in this work. We would also like to thank Megan Rigby for assistance with establishing the BRET measurements. We thank Ting-Yu Lin (UCSF) for providing cardiomyocyte isolation technical advice and Emanuela Zacco (UCSF-Laboratory for Cell Analysis) for technical support.

## Author contributions

Conceptualization: AC-N, AGS, PC

Data curation: AC-N

Formal Analysis: AC-N, MW, MS, DKS

Funding acquisition: AC-N, FM, AGS, PC

Investigation: AC-N, MW, RV, MS, AC, PC

Methodology: AC-N, MJS, DKS, AGS, PC

Resources: MRA, SM, MJS

Supervision: MJS, FM, AGS, PC

Visualization: AC-N, MW, MS

Writing – original draft: AC-N, DKS, AGS, PC

Writing – review & editing: all authors

## Data and materials availability

The RNA-Seq data from this manuscript have been deposited in NCBI’s Gene Expression Omnibus (*62*) and assigned the identifiers: GSE207187 and GSE207188. All other data needed to evaluate the conclusions in the paper are present in the paper or the Supplementary Materials.

NCBI GEO Private access:

Reviewer token (GSE207187): mfgfyyaifhmztmp

Reviewer token (GSE207188): yludoioydjmxjex

## Supplementary File

Fig. S1-S7

Tables S1-S3

## Supplementary Figures

**fig. S1.**
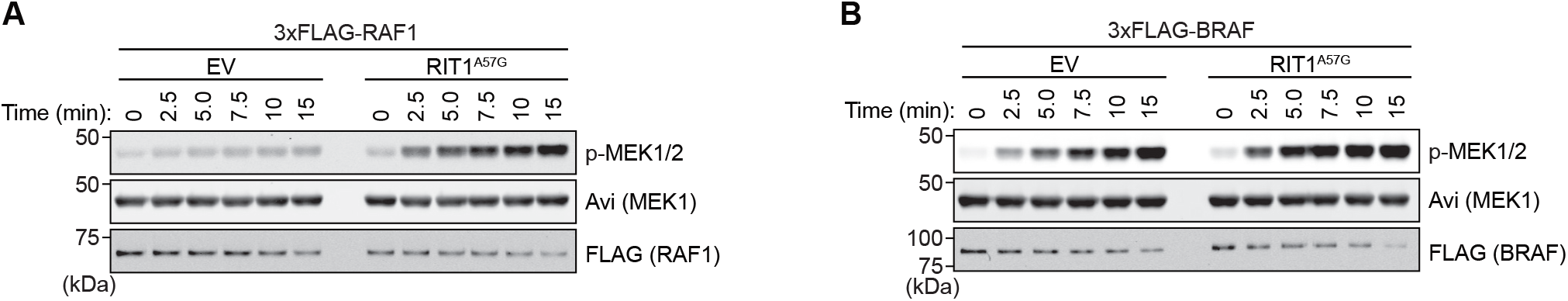
In vitro RAF kinase assay with RIT1^A57G^ co-expression. (**A, B**) Immunoblot analysis of kinase-dead MEK1 phosphorylation at indicated times by RAF1 (A) or BRAF (B) protein isolated from cells co-expressing RIT1^A57G^ or an empty vector (EV) control, in an in-vitro assay, see Methods.

**fig. S2.**
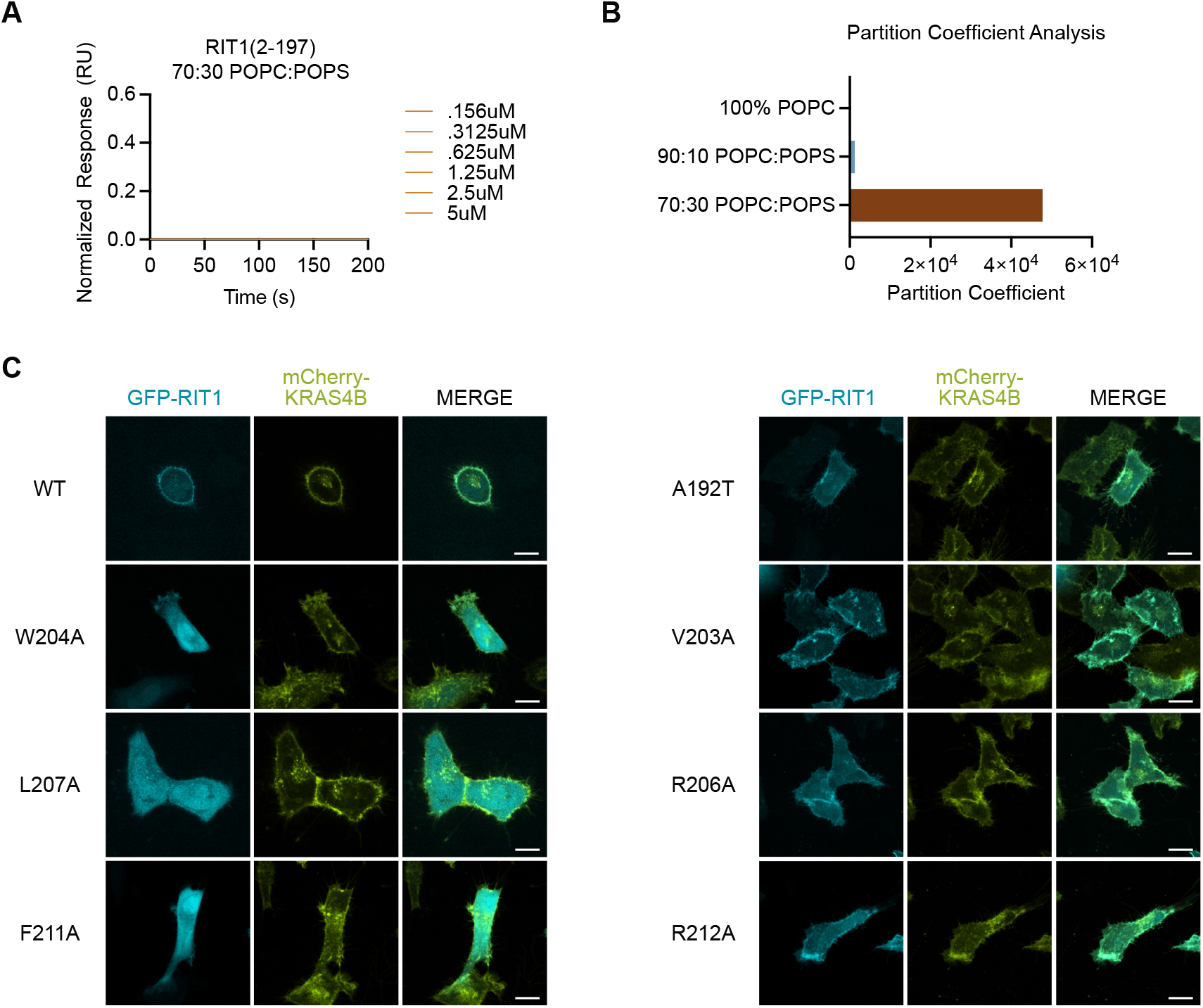
In vitro and in vivo analysis of RIT1-PM lipid associations. (**A**) SPR analysis shows no association between 30% POPS-containing liposomes and RIT1 protein (a.a. 2-197) with C-terminal deletion. (**B**) Partition coefficients derived from SPR affinity curves from Figure 1B show the relative binding affinity of RIT1 to liposomes of indicated composition. (**C**) Live-cell confocal images of HeLa cells transiently transfected with indicated GFP-RIT1 constructs. Stable expression of mCherry-KRAS4B was used as a plasma membrane marker. Representative images from one of three independent experiments (*n* = 3). Scale bar is 15 μm.

**fig. S3.**
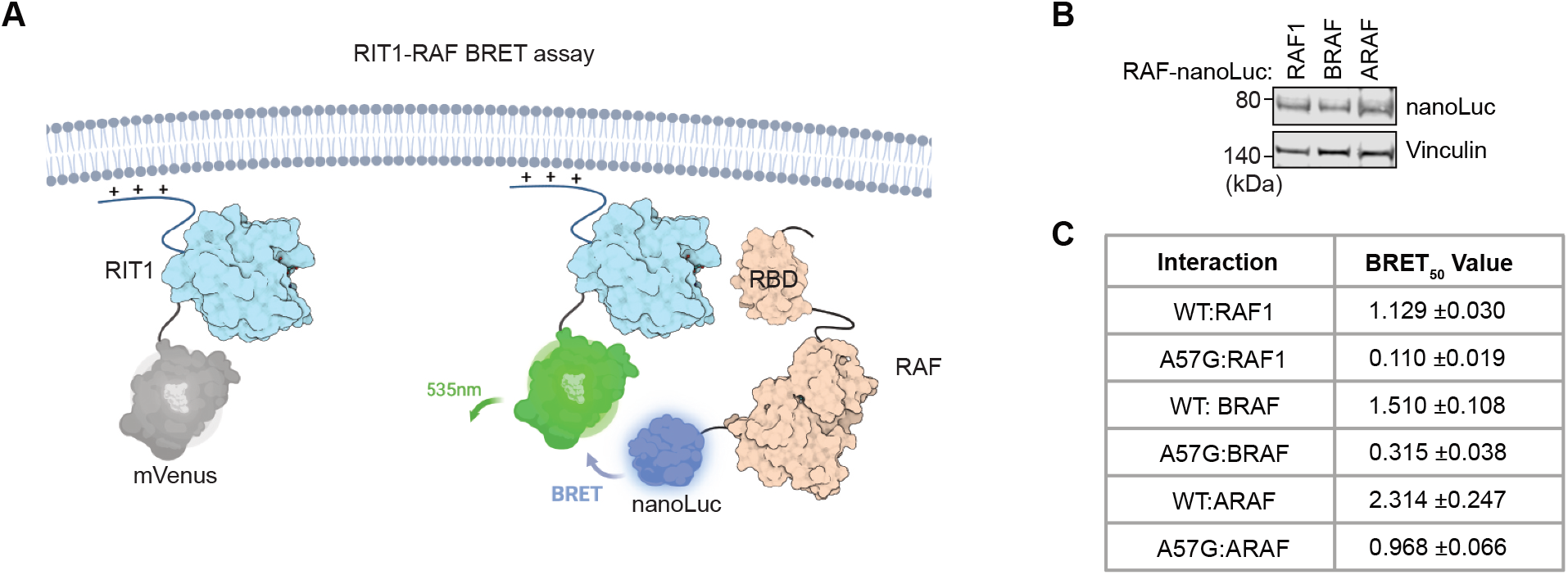
RIT1 BRET assay reveals preferential binding to RAF1 isoform. (**A**) Schema of BRET assay designed to detect in vivo RIT1-RAF binding. (**B**) RAF-nanoLuc construct transfection was optimized to achieve comparable RAF isoform expression levels. (**C**) BRET_50_ values calculated from saturation curves in Fig. 3B.

**fig. S4.**
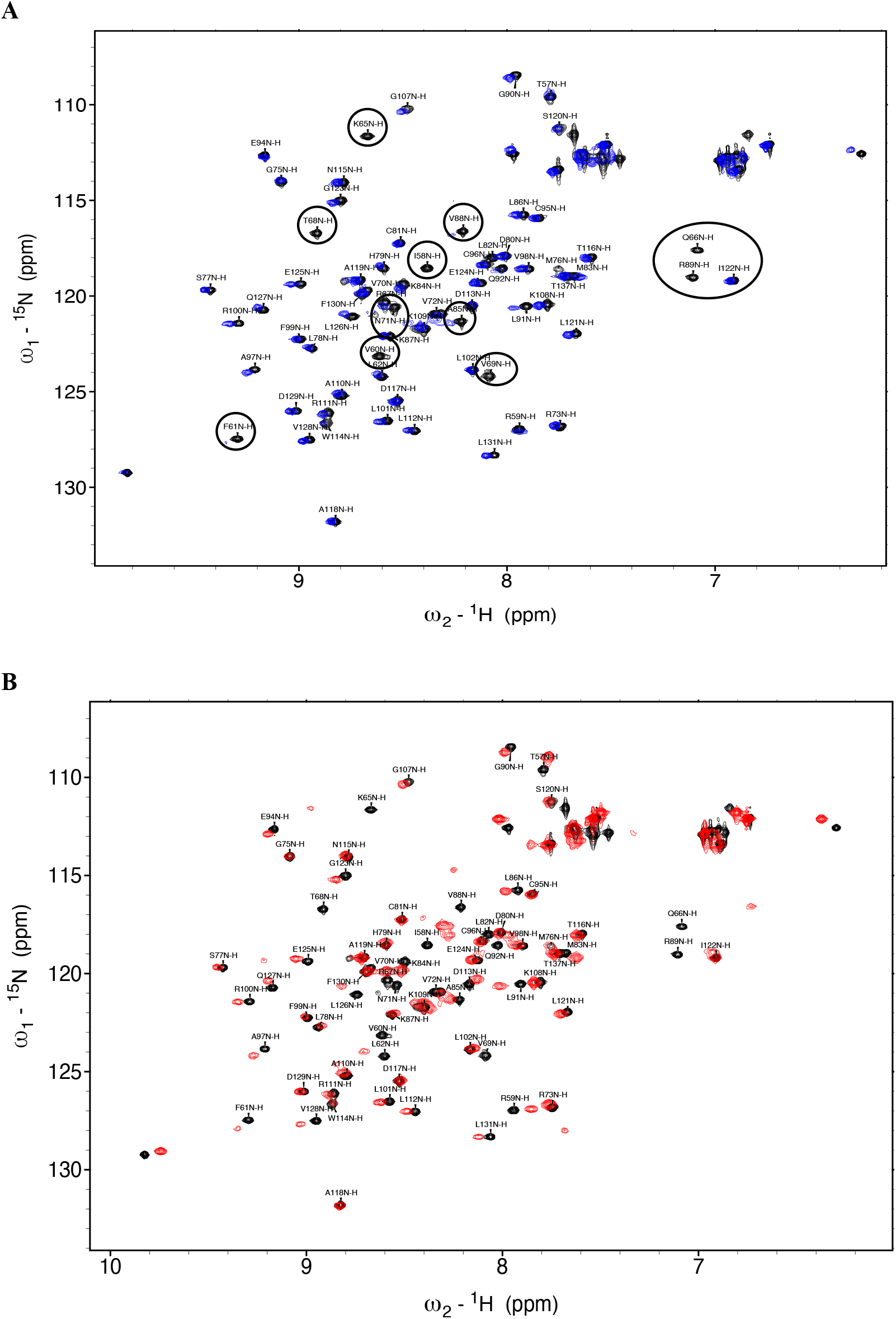
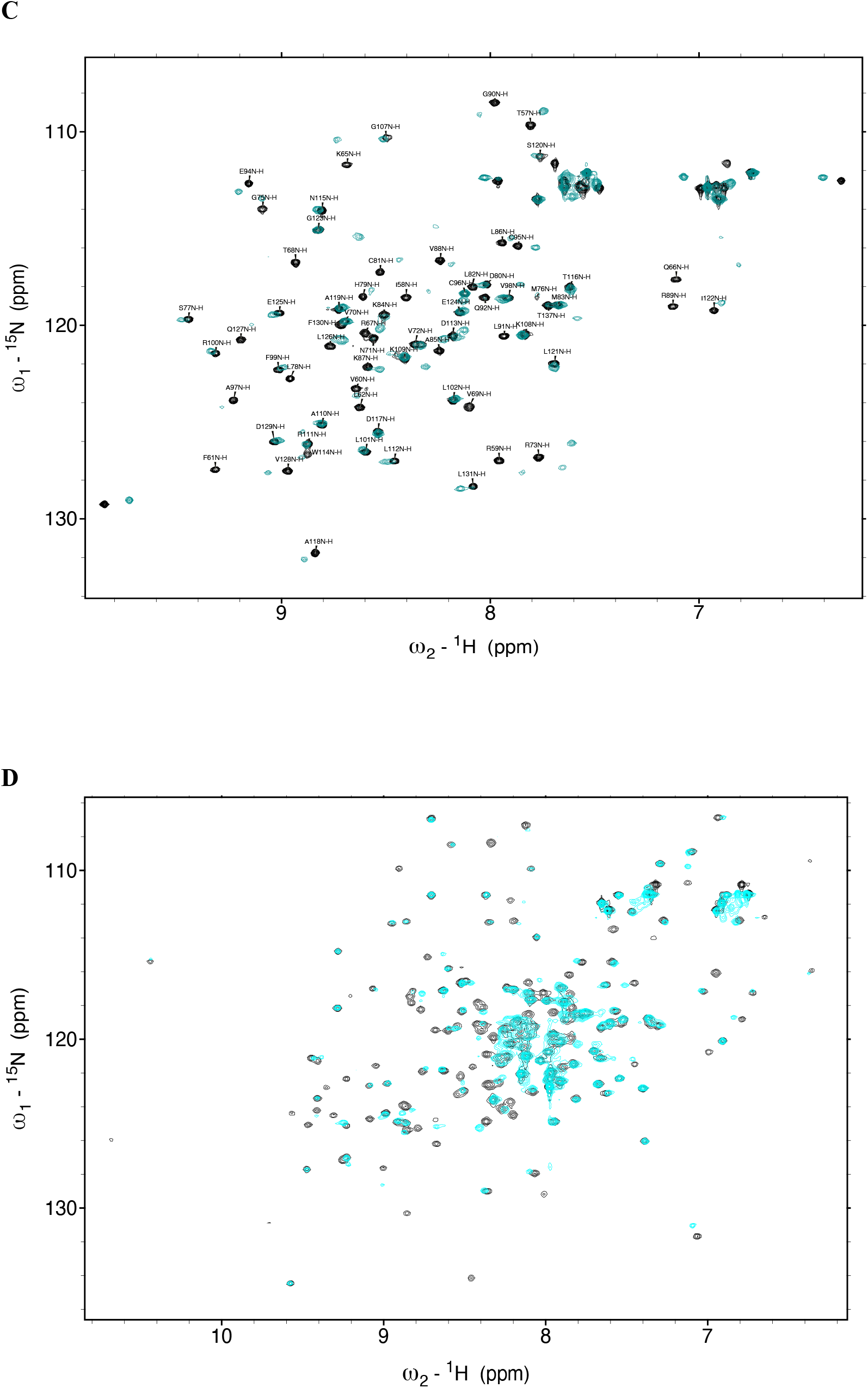
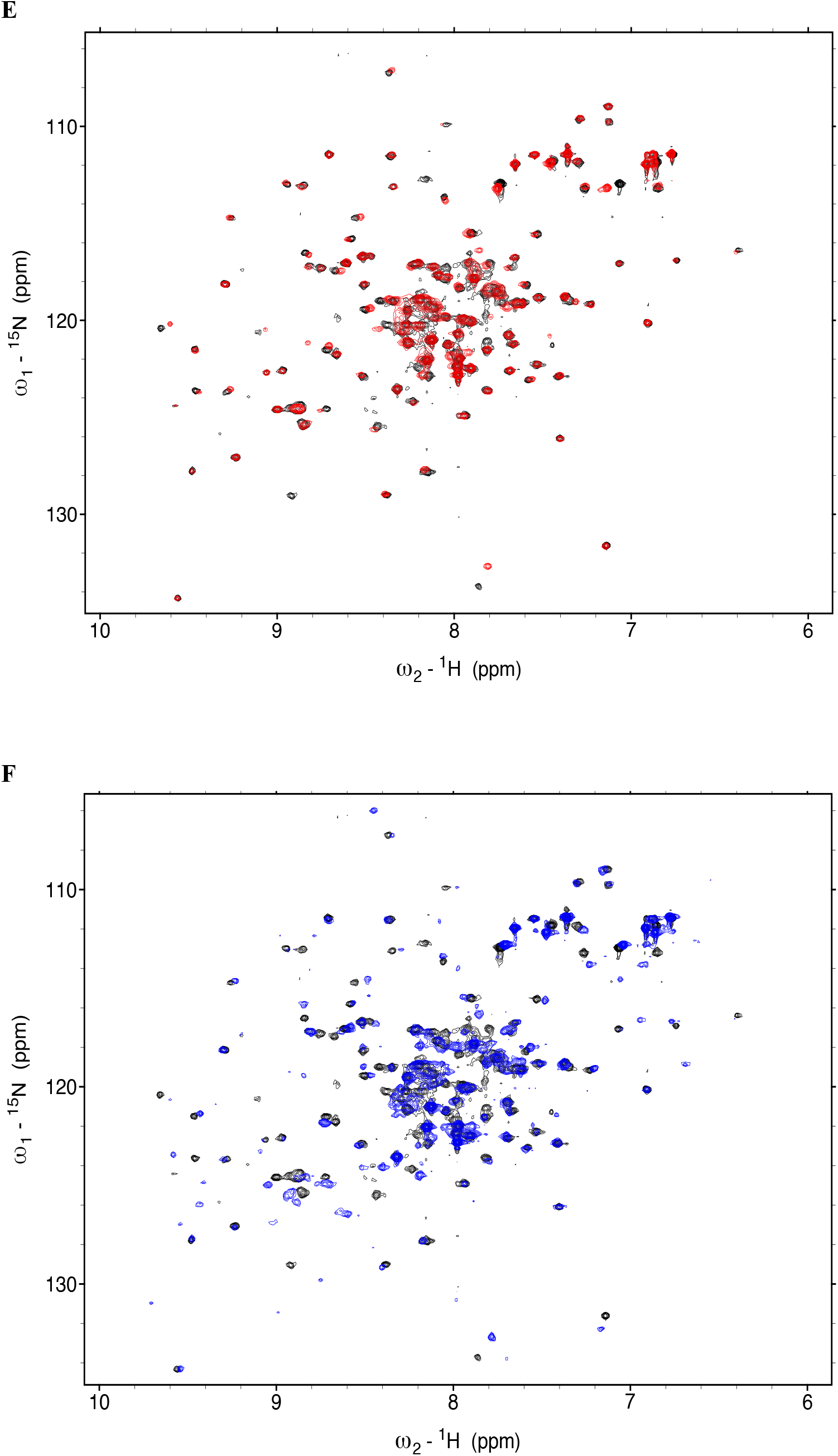
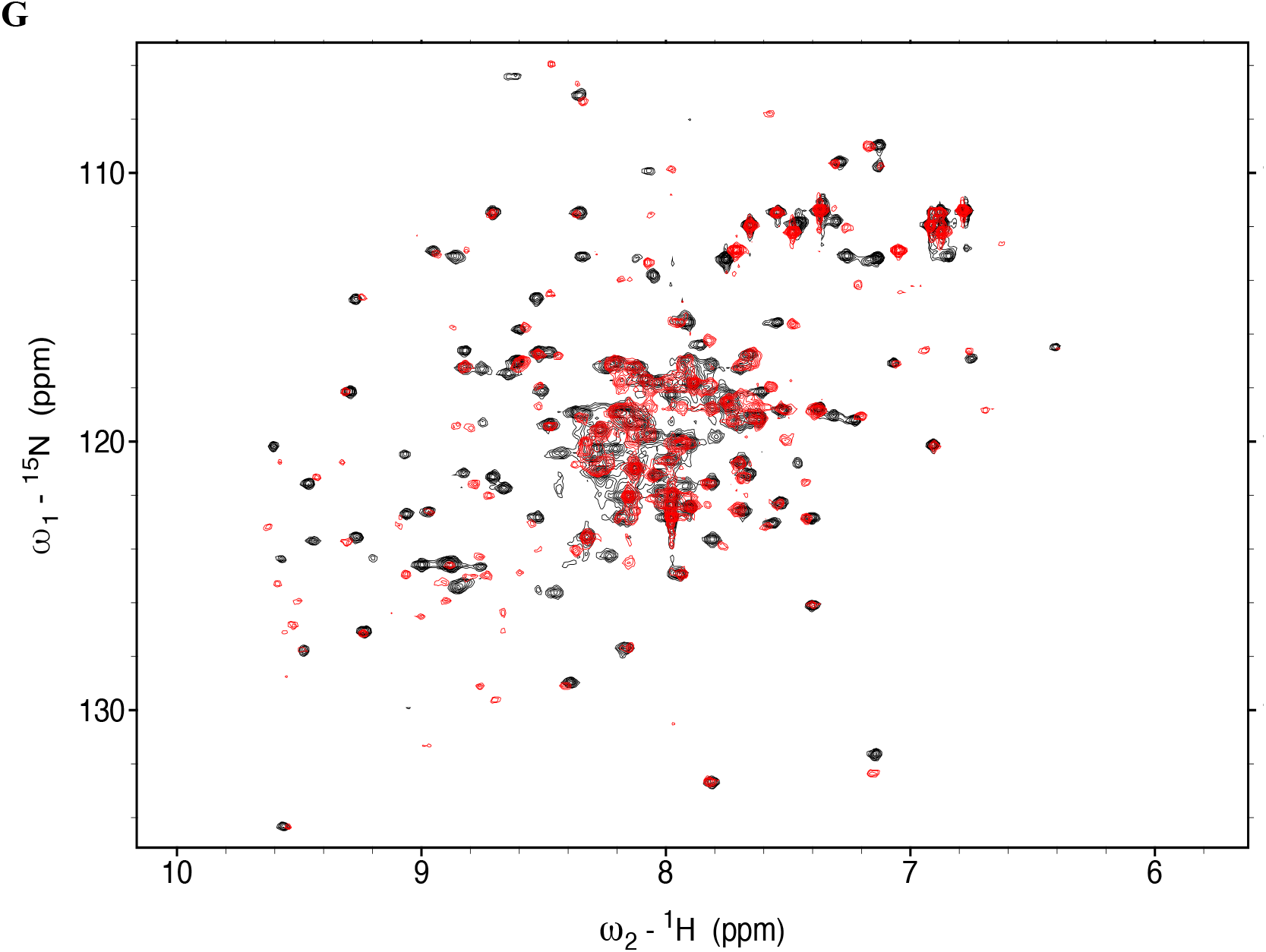
NMR spectra of RIT1-RAF1 RBD complexes. (**A**) Overlay ^15^N -^1^H HSQC spectra of ^15^N RBD (black) and ^15^N RBD in complex with unlabeled WT-RIT1-GppNHp (blue). (**B**) Overlay ^15^N -^1^H HSQC spectra of ^15^N RBD (black) and ^15^N RBD in complex with unlabeled RIT1^A57G^-GppNHp (red). (**C**) Overlay ^15^N -^1^H HSQC spectra of ^15^N RBD (black) and ^15^N RBD in complex with unlabeled KRAS-GppNHp (cyan). (**D**) Overlay ^15^N-^1^H HSQC spectra of ^15^N labeled WT-RIT1-GDP (black) with ^15^N labeled RIT1^A57G^-GDP (cyan). (**E**) Overlay ^15^N-^1^H HSQC spectra of ^15^N labeled WT-RIT1-GppNHp (black) with ^15^N labeled RIT1^A57G^-GppNHp (red). (**F**) Overlay ^15^N-^1^H HSQC spectra of ^15^N labeled RIT1-GppNHp in complex with RBD (blue) on ^15^N labeled RIT1-GppNHp (black). (**G**) Overlay ^15^N-^1^H HSQC spectra of ^15^N labeled RIT1^A57G^-GppNHp in complex with RBD (red) on ^15^N labeled RIT1^A57G^-GppNHp (black).

**fig. S5.**
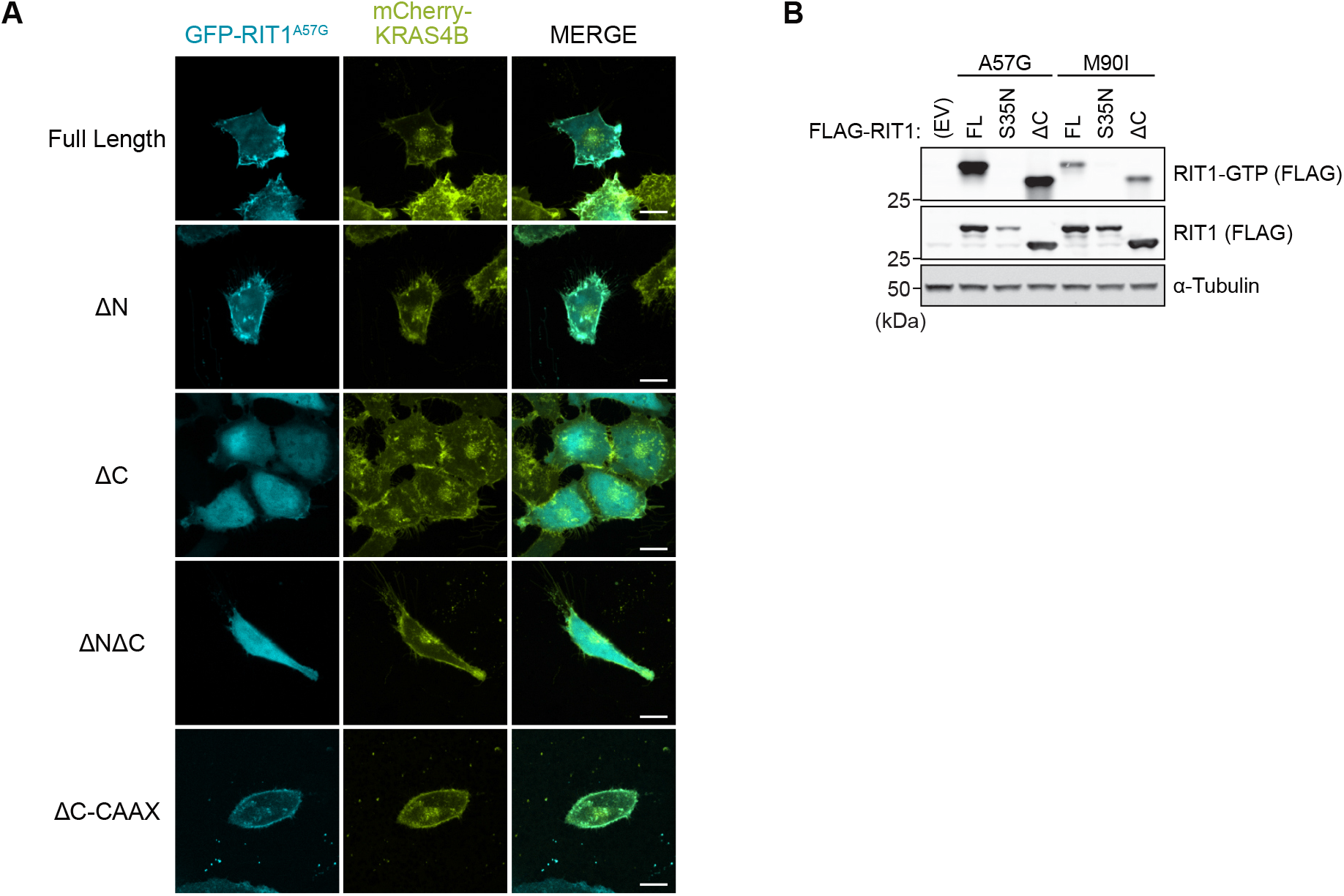
RIT1 C-terminus is essential for PM association but not GTP loading. (**A**) Live-cell confocal images of HeLa cells transiently transfected with indicated GFP-RIT1 constructs. Stable expression of mCherry-KRAS4B was used as a plasma membrane marker. Representative images from one of three independent experiments (*n* = 3). Scale bar is 15 μm. (**B**) Immunoblot analysis of indicated proteins from HEK-293 cells transiently transfected with indicated FLAG-tagged constructs and serum-starved for 16 h. GTP-bound RIT1 was precipitated with immobilized RAF1-RBD. One of two independent experiments is shown.

**fig. S6.**
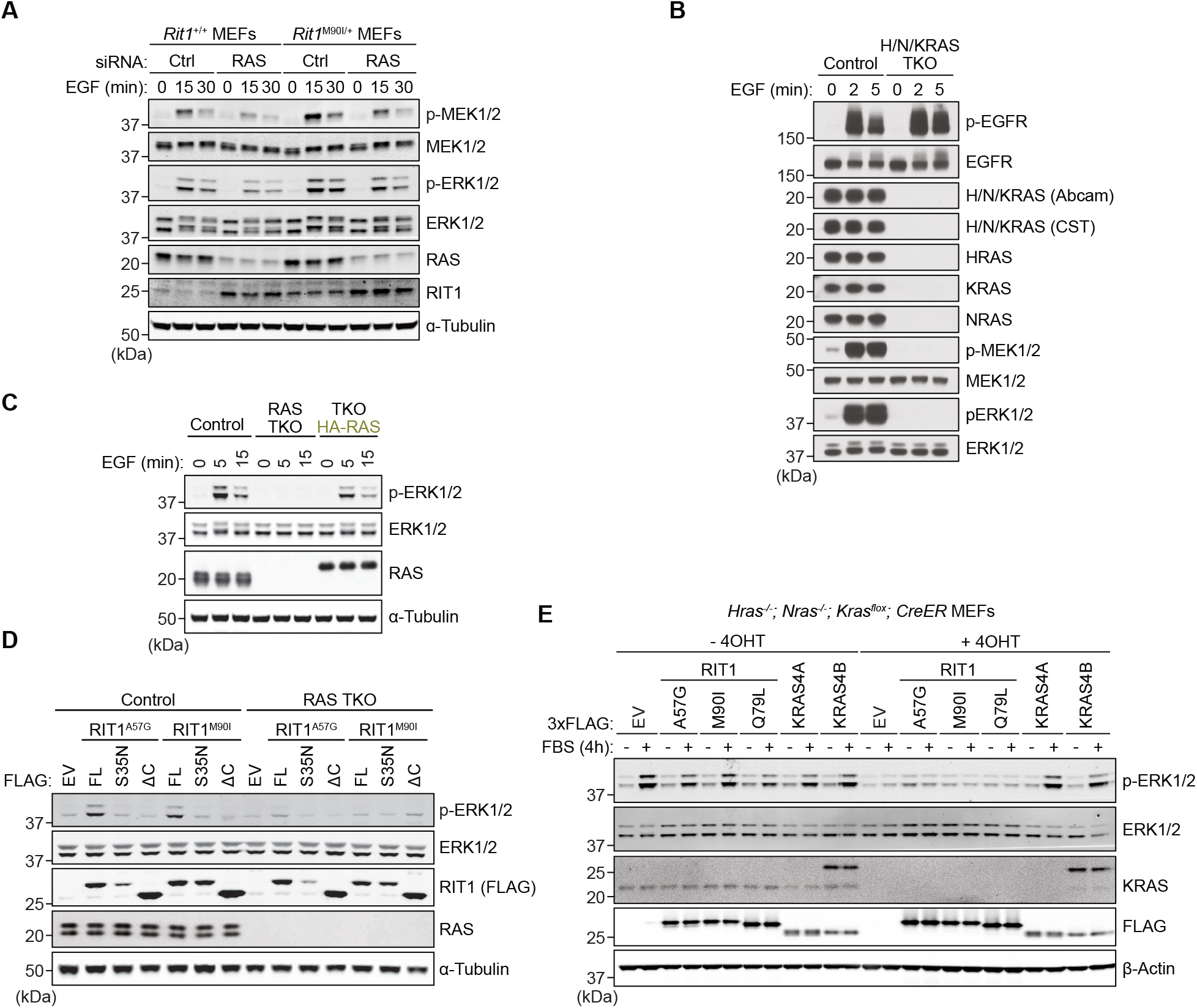
RIT1 protein stabilization fails to promote MAPK activation in the absence of RAS. (**A**) Immunoblot analysis of indicated proteins from primary MEFs that were serum-starved overnight and stimulated with 10 ng/ml EGF for indicated times, 72h post siRNA knockdown. One of three independent experiments is shown. (**B**) Immunoblot analysis of indicated proteins from Rasless (HRAS/NRAS/KRAS TKO) or control HEK-293 FlpIn cells serum-starved for 16 h and treated with 25 ng/ml EGF for indicated times. (**C**) Immunoblot analysis of indicated proteins from Rasless or control cells serum-starved for 16 h and treated with 10 ng/ml EGF for indicated times. Rasless cells were rescued with ectopic expression of HA-tagged HRAS, NRAS, KRAS4A, and KRAS4B (1:1:1:1 DNA ratio). One of two independent experiments is shown. (**D**) Immunoblot analysis of indicated proteins from Rasless or control cells transiently transfected with indicated FLAG-tagged RIT1 constructs or an empty vector (EV) control and serum-starved for 16 h. FL, full length. One of three independent experiments is shown. (**E**) Immunoblot analysis of indicated proteins from control (−4OHT) and Rasless (+4OHT) MEFs stably expressing indicated constructs, serum-starved overnight and stimulated with or without 10% FBS for 4 h. One of three independent experiments is shown.

**fig. S7.**
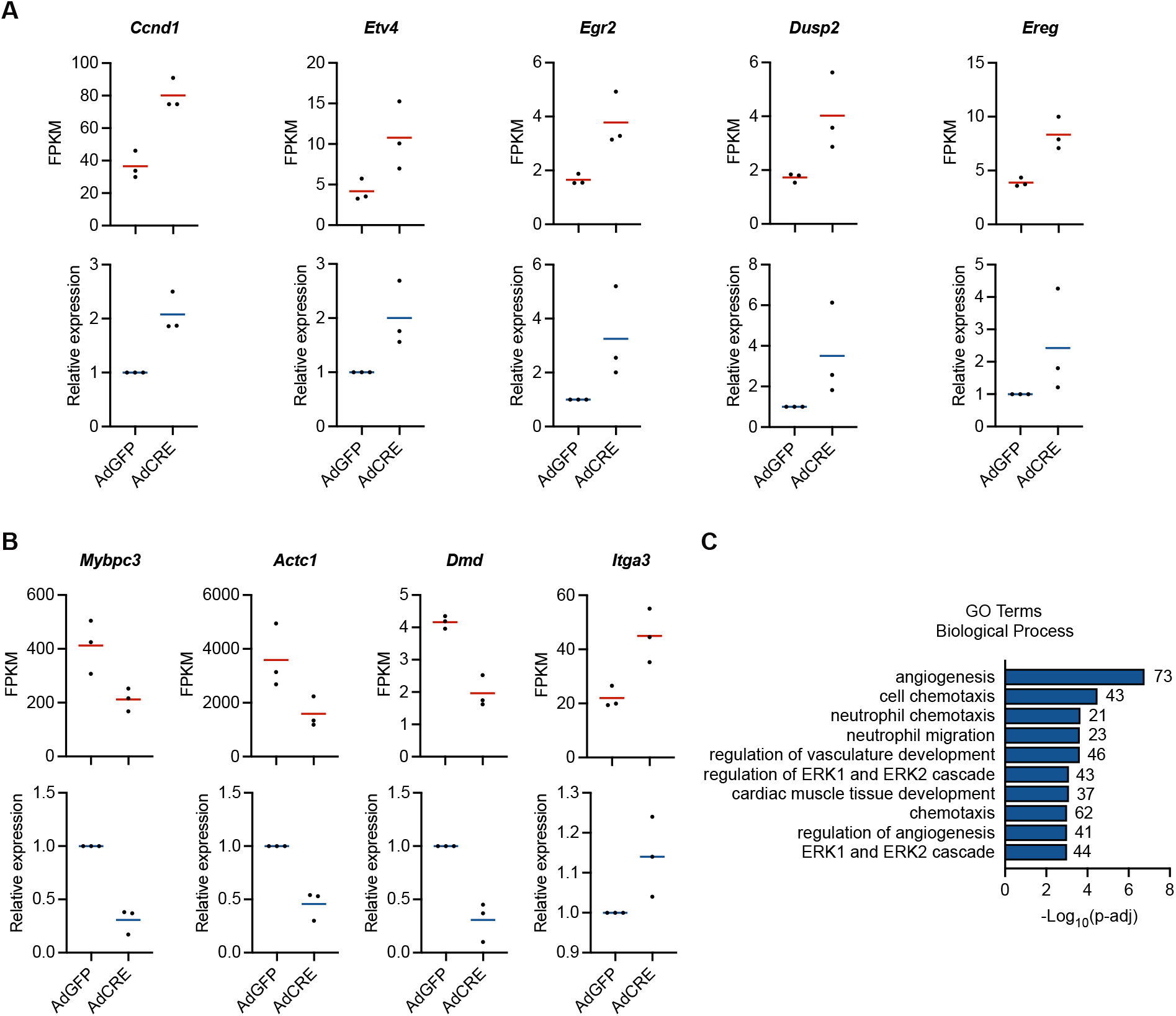
Analysis of gene expression elicited by RIT1^M90I^ expression or MEK inhibition. (**A, B**) Gene expression levels of indicated MAPK-regulated genes (A) or cardiomyopathy-associated genes (B) from primary *Rit1*^*LoxP-M90I*^ neonatal cardiomyocytes treated with adenovirus encoding Cre recombinase (AdCre) or GFP (AdGFP). Top panels display normalized mRNA transcript levels (FPKM) from RNA-seq transcriptomic profiling. Bottom panels display relative mRNA expression from an independent set of biological replicates by RT-qPCR. (**C**) Gene ontology (GO) enrichment analysis of genes downregulated in MEKi-treated hearts (MEKi vs. vehicle control) isolated from the murine preclinical trial described in Fig. 8D.

## Supplementary Tables

**Table S1.** Broadened residues (blue) and residues above 1.5σ reported in CSP plots.

| <sup>15</sup>N RBD<br>+ RIT1 | <sup>15</sup>N RBD<br>+ RIT1<sup>A57G</sup> | <sup>15</sup>N RBD<br>+ KRAS | <sup>15</sup>N RIT1<br>+ RBD | <sup>15</sup>N RIT1<sup>A57G</sup><br>+ RBD | <sup>15</sup>N KRAS<br>+ RBD |
| --- | --- | --- | --- | --- | --- |
| T57 | T57 | T57 | R20 | R20 | Y4 |
| I58 | I58 | I58 | E21 | E21 | L6 |
| V60 | R59 | R59 | M26 | M26 | V8 |
| F61 | V60 | V60 | L27 | L27 | V9 |
| L62 | F61 | F61 | A29 | G30 | G15 |
| K65 | L62 | L62 | H44 | M37 | S17 |
| Q66 | K65 | K65 | A57 | S43 | Q25 |
| T68 | Q66 | Q66 | Y58 | H44 | H27 |
| V69 | R67 | R67 | I62 | K59 | F28 |
| N71 | T68 | T68 | A69 | I62 | V45 |
| V88 | V69 | V69 | N70 | A69 | D47 |
| R89 | V70 | V70 | L71 | L71 | L56 |
| Q92 | N71 | R73 | Q79 | Q79 | T74 |
| A97 | L82 | G75 | M85 | A84 | G75 |
| L126 | M83 | L78 | R86 | M85 | I84 |
| Q127 | K84 | H79 | E94 | R86 | T87 |
|  | V88 | C81 | G95 | E94 | E91 |
|  | R89 | A85 | F96 | G95 | Y96 |
|  | G90 | K87 | S101 | F96 | V103 |
|  | L91 | V88 | E110 | Y100 | D108 |
|  | Q92 | R89 | R112 | S101 | R123 |
|  | E94 | G90 | V121 | V111 | Y137 |
|  | A97 | L91 | R123 | R112 | R149 |
|  | D113 | Q92 | G133 | Y119 | F156 |
|  | E125 | E94 | S164 | R123 | Y157 |
|  | L126 | A97 | A166 | V132 |  |
|  | Q127 | A118 | Y170 | G133 |  |
|  | V128 | I122 | D172 | S164 |  |
|  | D129 | L126 | F175 | R168 |  |
|  |  | Q127 | R180 | Y170 |  |
|  |  | V128 | L191 | D172 |  |
|  |  | L131 |  | R180 |  |

**Table S2.**
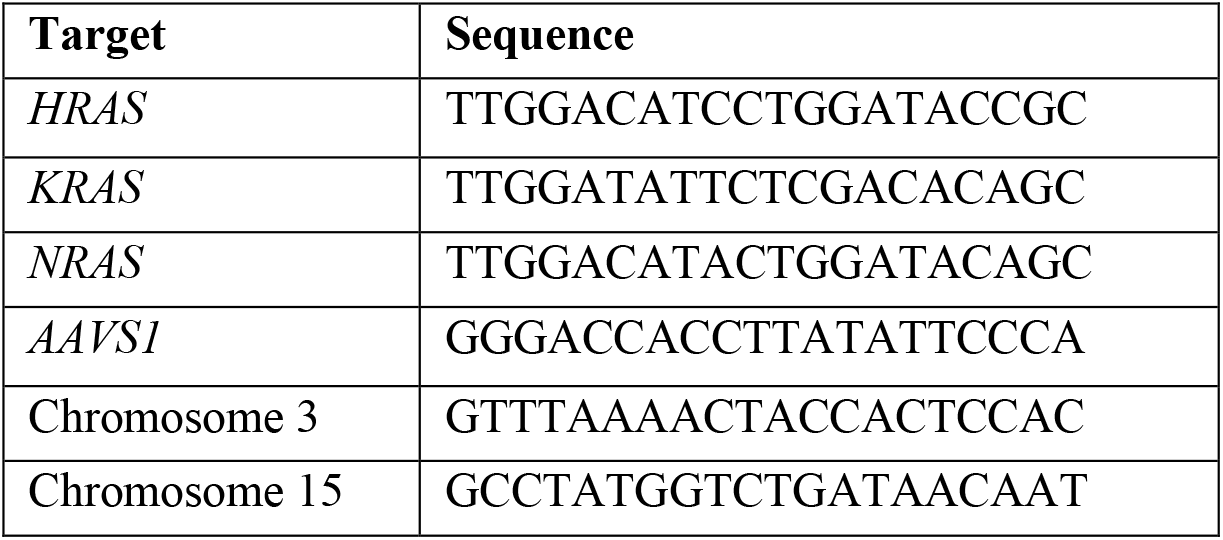
gRNA targeting sequences.

**Table S3.**
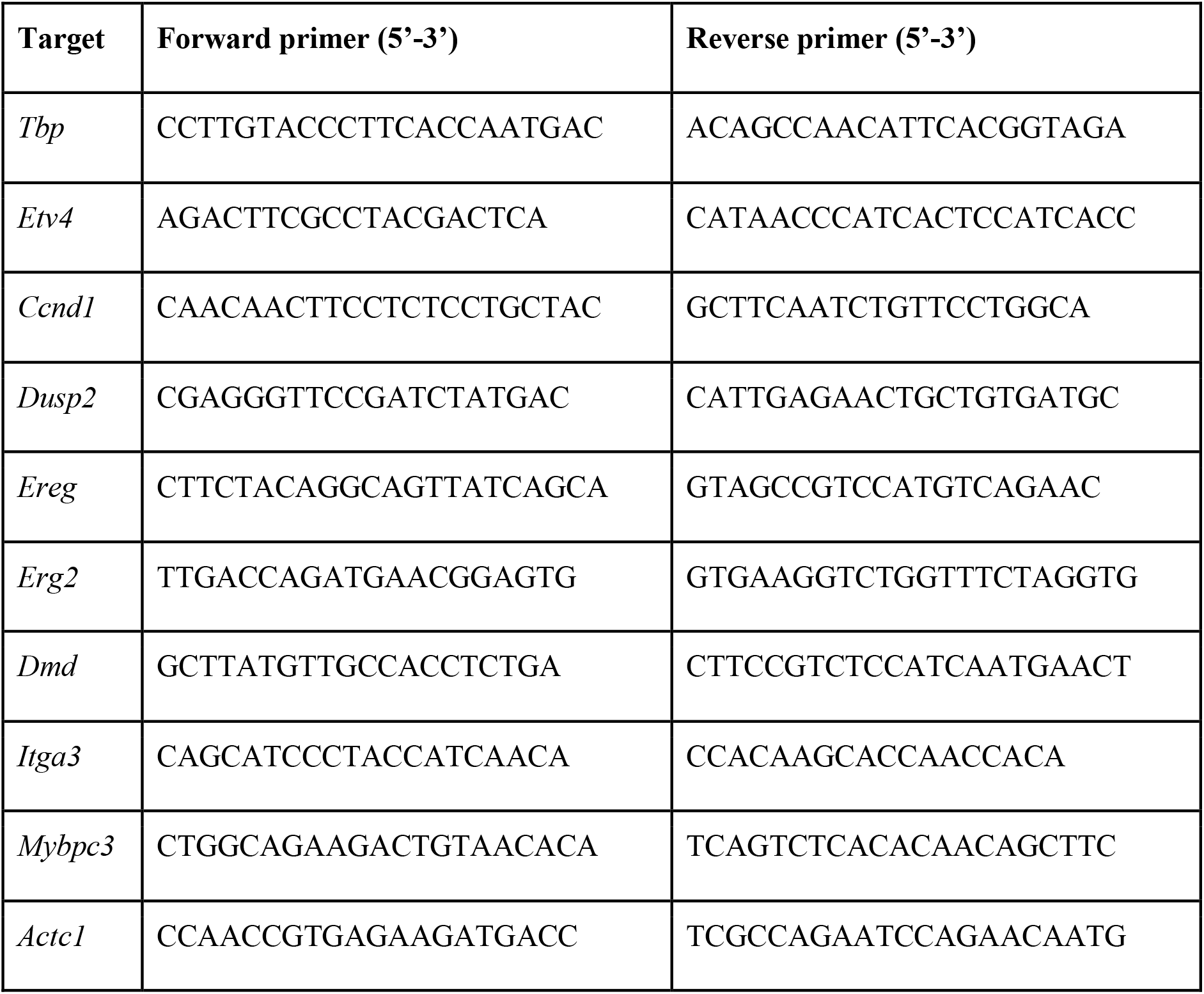
qPCR primers.

## References

1. P. Castel, K. A. Rauen, F. McCormick, The duality of human oncoproteins: drivers of cancer and congenital disorders. Nat Rev Cancer. 20, 383–397 (2020).

2. D. K. Simanshu, D. V. Nissley, F. McCormick, RAS Proteins and Their Regulators in Human Disease. Cell. 170, 17–33 (2017).

3. S. J. Leevers, H. F. Paterson, C. J. Marshall, Requirement for Ras in Raf activation is overcome by targeting Raf to the plasma membrane. Nature. 369, 411–414 (1994).

4. D. Stokoe, S. G. Macdonald, K. Cadwallader, M. Symons, J. F. Hancock, Activation of Raf as a result of recruitment to the plasma membrane. Science. 264, 1463–1467 (1994).

5. K. A. Rauen, The RASopathies. Annu Rev Genomics Hum Genet. 14, 355–369 (2013).

6. A. H. Berger, M. Imielinski, F. Duke, J. Wala, N. Kaplan, G.-X. Shi, D. A. Andres, M. Meyerson, Oncogenic RIT1 mutations in lung adenocarcinoma. Oncogene. 33, 4418–4423 (2014).

7. I. Gómez-Seguí, H. Makishima, A. Jerez, K. Yoshida, B. Przychodzen, S. Miyano, Y. Shiraishi, H. D. Husseinzadeh, K. Guinta, M. Clemente, N. Hosono, M. A. McDevitt, A. R. Moliterno, M. A. Sekeres, S. Ogawa, J. P. Maciejewski, Novel recurrent mutations in the RAS-like GTP-binding gene RIT1 in myeloid malignancies. Leukemia. 27, 1943–1946 (2013).

8. R. Van, A. Cuevas-Navarro, P. Castel, F. McCormick, The molecular functions of RIT1 and its contribution to human disease. Biochem J. 477, 2755–2770 (2020).

9. M. Yaoita, T. Niihori, S. Mizuno, N. Okamoto, S. Hayashi, A. Watanabe, M. Yokozawa, H. Suzumura, A. Nakahara, Y. Nakano, T. Hokosaki, A. Ohmori, H. Sawada, O. Migita, A. Mima, P. Lapunzina, F. Santos-Simarro, S. García-Miñaúr, T. Ogata, H. Kawame, K. Kurosawa, H. Ohashi, S. -Inoue, Y. Matsubara, S. Kure, Y. Aoki, Spectrum of mutations and genotype-phenotype analysis in Noonan syndrome patients with RIT1 mutations. Hum Genet. 135, 209–222 (2016).

10. Y. Aoki, T. Niihori, T. Banjo, N. Okamoto, S. Mizuno, K. Kurosawa, T. Ogata, F. Takada, M. Yano, T. Ando, T. Hoshika, C. Barnett, H. Ohashi, H. Kawame, T. Hasegawa, T. Okutani, T. Nagashima, S. Hasegawa, R. Funayama, T. Nagashima, K. Nakayama, S.-I. Inoue, Y. Watanabe, T. Ogura, Y. Matsubara, Gain-of-function mutations in RIT1 cause Noonan syndrome, a RAS/MAPK pathway syndrome. Am J Hum Genet. 93, 173–180 (2013).

11. M. Koenighofer, C. Y. Hung, J. L. McCauley, J. Dallman, E. J. Back, I. Mihalek, K. W. Gripp, K. Sol-Church, P. Rusconi, Z. Zhang, G.-X. Shi, D. A. Andres, O. A. Bodamer, Mutations in RIT1 cause Noonan syndrome - additional functional evidence and expanding the clinical phenotype. Clin Genet. 89, 359–366 (2016).

12. A. E. Roberts, J. E. Allanson, M. Tartaglia, B. D. Gelb, Noonan syndrome. Lancet. 381, 333–342 (2013).

13. P. Castel, A. Cheng, A. Cuevas-Navarro, D. B. Everman, A. G. Papageorge, D. K. Simanshu, A. Tankka, J. Galeas, A. Urisman, F. McCormick, RIT1 oncoproteins escape LZTR1-mediated proteolysis. Science. 363, 1226–1230 (2019).

14. S. Takahara, S.-I. Inoue, S. Miyagawa-Tomita, K. Matsuura, Y. Nakashima, T. Niihori, Y. Matsubara, Y. Saiki, Y. Aoki, New Noonan syndrome model mice with RIT1 mutation exhibit cardiac hypertrophy and susceptibility to β-adrenergic stimulation-induced cardiac fibrosis. EBioMedicine. 42, 43–53 (2019).

15. A. Cuevas-Navarro, L. Rodriguez-Muñoz, J. Grego-Bessa, A. Cheng, K. A. Rauen, A. Urisman, F. McCormick, G. Jimenez, P. Castel, Cross-species analysis of LZTR1 loss-of-function mutants demonstrates dependency to RIT1 orthologs. Elife. 11, e76495 (2022).

16. Y. Wang, J. Zhang, P. Zhang, Z. Zhao, Q. Huang, D. Yun, J. Chen, H. Chen, C. Wang, D. Lu, bioRxiv, in press, doi:10.1101/2020.03.14.989954.

17. G.-X. Shi, D. A. Andres, Rit contributes to nerve growth factor-induced neuronal differentiation via activation of B-Raf-extracellular signal-regulated kinase and p38 mitogen-activated protein kinase cascades. Mol Cell Biol. 25, 830–846 (2005).

18. S.-B. Peng, J. R. Henry, M. D. Kaufman, W.-P. Lu, B. D. Smith, S. Vogeti, T. J. Rutkoski, S. Wise, L. Chun, Y. Zhang, R. D. Van Horn, T. Yin, X. Zhang, V. Yadav, S.-H. Chen, X. Gong, X. Ma, Y. Webster, S. Buchanan, I. Mochalkin, L. Huber, L. Kays, G. P. Donoho, J. Walgren, D. McCann, P. Patel, I. Conti, G. D. Plowman, J. J. Starling, D. L. Flynn, Inhibition of RAF Isoforms and Active Dimers by LY3009120 Leads to Anti-tumor Activities in RAS or BRAF Mutant Cancers. Cancer Cell. 28, 384–398 (2015).

19. R. Marais, Y. Light, H. F. Paterson, C. S. Mason, C. J. Marshall, Differential regulation of Raf-1, A-Raf, and B-Raf by oncogenic ras and tyrosine kinases. J Biol Chem. 272, 4378–4383 (1997).

20. C. S. Mason, C. J. Springer, R. G. Cooper, G. Superti-Furga, C. J. Marshall, R. Marais, Serine and tyrosine phosphorylations cooperate in Raf-1, but not B-Raf activation. EMBO J. 18, 2137–2148 (1999).

21. T. N. Figueira, J. M. Freire, C. Cunha-Santos, M. Heras, J. Gonçalves, A. Moscona, M. Porotto, A. Salomé Veiga, M. A. R. B. Castanho, Quantitative analysis of molecular partition towards lipid membranes using surface plasmon resonance. Sci Rep. 7, 45647 (2017).

22. A. D. Migliori, L. A. Patel, C. Neale, The RIT1 C-terminus associates with lipid bilayers via charge complementarity. Comput Biol Chem. 91, 107437 (2021).

23. W. D. Heo, T. Inoue, W. S. Park, M. L. Kim, B. O. Park, T. J. Wandless, T. Meyer, PI(3,4,5)P3 and PI(4,5)P2 lipids target proteins with polybasic clusters to the plasma membrane. Science. 314, 1458– 1461 (2006).

24. E. M. Terrell, D. E. Durrant, D. A. Ritt, N. E. Sealover, E. Sheffels, R. Spencer-Smith, D. Esposito, Y. Zhou, J. F. Hancock, R. L. Kortum, D. K. Morrison, Distinct Binding Preferences between Ras and Raf Family Members and the Impact on Oncogenic Ras Signaling. Mol Cell. 76, 872-884.e5 (2019).

25. T. H. Tran, A. H. Chan, L. C. Young, L. Bindu, C. Neale, S. Messing, S. Dharmaiah, T. Taylor, J.-P. Denson, D. Esposito, D. V. Nissley, A. G. Stephen, F. McCormick, D. K. Simanshu, KRAS interaction with RAF1 RAS-binding domain and cysteine-rich domain provides insights into RAS-mediated RAF activation. Nat Commun. 12, 1176 (2021).

26. P. Rodriguez-Viciana, P. H. Warne, A. Khwaja, B. M. Marte, D. Pappin, P. Das, M. D. Waterfield, A. Ridley, J. Downward, Role of phosphoinositide 3-OH kinase in cell transformation and control of the actin cytoskeleton by Ras. Cell. 89, 457–467 (1997).

27. H. Nakhaeizadeh, E. Amin, S. Nakhaei-Rad, R. Dvorsky, M. R. Ahmadian, The RAS-Effector Interface: Isoform-Specific Differences in the Effector Binding Regions. PLoS One. 11, e0167145 (2016).

28. G. N. Ramachandran, C. Ramakrishnan, V. Sasisekharan, Stereochemistry of polypeptide chain configurations. J Mol Biol. 7, 95–99 (1963).

29. N. Bhattacharjee, P. Biswas, Position-specific propensities of amino acids in the β-strand. BMC Struct Biol. 10, 29 (2010).

30. E. M. Terrell, D. K. Morrison, Ras-Mediated Activation of the Raf Family Kinases. Cold Spring Harb Perspect Med. 9, a033746 (2019).

31. A. Cuevas-Navarro, R. Van, A. Cheng, A. Urisman, P. Castel, F. McCormick, The RAS GTPase RIT1 compromises mitotic fidelity through spindle assembly checkpoint suppression. Curr Biol. 31, 3915-3924.e9 (2021).

32. L. A. Feig, G. M. Cooper, Inhibition of NIH 3T3 cell proliferation by a mutant ras protein with preferential affinity for GDP. Mol Cell Biol. 8, 3235–3243 (1988).

33. U. Meyer Zum Büschenfelde, L. I. Brandenstein, L. von Elsner, K. Flato, T. Holling, M. Zenker, G. Rosenberger, K. Kutsche, RIT1 controls actin dynamics via complex formation with RAC1/CDC42 and PAK1. PLoS Genet. 14, e1007370 (2018).

34. P. Rodriguez-Viciana, C. Sabatier, F. McCormick, Signaling specificity by Ras family GTPases is determined by the full spectrum of effectors they regulate. Mol Cell Biol. 24, 4943–4954 (2004).

35. M. Drosten, A. Dhawahir, E. Y. M. Sum, J. Urosevic, C. G. Lechuga, L. M. Esteban, E. Castellano, C. Guerra, E. Santos, M. Barbacid, Genetic analysis of Ras signalling pathways in cell proliferation, migration and survival. EMBO J. 29, 1091–1104 (2010).

36. L. Sun, S. Xi, Z. Zhou, F. Zhang, P. Hu, Y. Cui, S. Wu, Y. Wang, S. Wu, Y. Wang, Y. Du, J. Zheng, H. Yang, M. Chen, Q. Yan, D. Yu, C. Shi, Y. Zhang, D. Xie, X.-Y. Guan, Y. Li, Elevated expression of RIT1 hyperactivates RAS/MAPK signal and sensitizes hepatocellular carcinoma to combined treatment with sorafenib and AKT inhibitor. Oncogene. 41, 732–744 (2022).

37. F. Uhlitz, A. Sieber, E. Wyler, R. Fritsche-Guenther, J. Meisig, M. Landthaler, B. Klinger, N. Blüthgen, An immediate-late gene expression module decodes ERK signal duration. Mol Syst Biol. 13, 928 (2017).

38. F. Sedaghat-Hamedani, E. Kayvanpour, O. F. Tugrul, A. Lai, A. Amr, J. Haas, T. Proctor, P. Ehlermann, K. Jensen, H. A. Katus, B. Meder, Clinical outcomes associated with sarcomere mutations in hypertrophic cardiomyopathy: a meta-analysis on 7675 individuals. Clin Res Cardiol. 107, 30–41 (2018).

39. A. J. Marian, E. Braunwald, Hypertrophic Cardiomyopathy: Genetics, Pathogenesis, Clinical Manifestations, Diagnosis, and Therapy. Circ Res. 121, 749–770 (2017).

40. C. H. Lee, N. G. Della, C. E. Chew, D. J. Zack, Rin, a neuron-specific and calmodulin-binding small G-protein, and Rit define a novel subfamily of ras proteins. J Neurosci. 16, 6784–6794 (1996).

41. Y. Aoki, T. Niihori, S. Inoue, Y. Matsubara, Recent advances in RASopathies. J Hum Genet. 61, 33– 39 (2016).

42. Z. Fang, C. B. Marshall, J. C. Yin, M. T. Mazhab-Jafari, G. M. C. Gasmi-Seabrook, M. J. Smith, T. Nishikawa, Y. Xu, B. G. Neel, M. Ikura, Biochemical Classification of Disease-associated Mutants of RAS-like Protein Expressed in Many Tissues (RIT1). J Biol Chem. 291, 15641–15652 (2016).

43. G. Andelfinger, C. Marquis, M.-J. Raboisson, Y. Théoret, S. Waldmüller, G. Wiegand, B. D. Gelb, M. Zenker, M.-A. Delrue, M. Hofbeck, Hypertrophic Cardiomyopathy in Noonan Syndrome Treated by MEK-Inhibition. J Am Coll Cardiol. 73, 2237–2239 (2019).

44. A. Leegaard, P. A. Gregersen, T. Ø. Nielsen, J. V. Bjerre, M. M. Handrup, Succesful MEK-inhibition of severe hypertrophic cardiomyopathy in RIT1-related Noonan Syndrome. Eur J Med Genet. 65, 104630 (2022).

45. J. R. Infante, L. A. Fecher, G. S. Falchook, S. Nallapareddy, M. S. Gordon, C. Becerra, D. J. DeMarini, D. S. Cox, Y. Xu, S. R. Morris, V. G. R. Peddareddigari, N. T. Le, L. Hart, J. C. Bendell, G. Eckhardt, R. Kurzrock, K. Flaherty, H. A. Burris, W. A. Messersmith, Safety, pharmacokinetic, pharmacodynamic, and efficacy data for the oral MEK inhibitor trametinib: a phase 1 dose-escalation trial. Lancet Oncol. 13, 773–781 (2012).

46. K. T. Flaherty, C. Robert, P. Hersey, P. Nathan, C. Garbe, M. Milhem, L. V. Demidov, J. C. Hassel, P. Rutkowski, P. Mohr, R. Dummer, U. Trefzer, J. M. G. Larkin, J. Utikal, B. Dreno, M. Nyakas, M. R. Middleton, J. C. Becker, M. Casey, L. J. Sherman, F. S. Wu, D. Ouellet, A.-M. Martin, K. Patel, D. Schadendorf, METRIC Study Group, Improved survival with MEK inhibition in BRAF-mutated melanoma. N Engl J Med. 367, 107–114 (2012).

47. A. Vichas, A. K. Riley, N. T. Nkinsi, S. Kamlapurkar, P. C. R. Parrish, A. Lo, F. Duke, J. Chen, I. Fung, J. Watson, M. Rees, A. M. Gabel, J. D. Thomas, R. K. Bradley, J. K. Lee, E. M. Hatch, M. K. Baine, N. Rekhtman, M. Ladanyi, F. Piccioni, A. H. Berger, Integrative oncogene-dependency mapping identifies RIT1 vulnerabilities and synergies in lung cancer. Nat Commun. 12, 4789 (2021).

48. W. Wei, M. J. Geer, X. Guo, I. Dolgalev, N. E. Sanjana, B. G. Neel, bioRxiv, in press, doi:10.1101/2022.08.26.505487.

49. S. Chen, R. S. Vedula, A. Cuevas-Navarro, B. Lu, S. J. Hogg, E. Wang, S. Benbarche, K. Knorr, W. J. Kim, R. F. Stanley, H. Cho, C. Erickson, M. Singer, D. Cui, S. Tittley, B. H. Durham, T. S. Pavletich, E. Fiala, M. F. Walsh, D. Inoue, S. Monette, J. Taylor, N. Rosen, F. McCormick, R. C. Lindsley, P. Castel, O. Abdel-Wahab, Impaired Proteolysis of Noncanonical RAS Proteins Drives Clonal Hematopoietic Transformation. Cancer Discovery, OF1–OF20 (2022).

50. L. C. Young, N. Hartig, I. Boned Del Río, S. Sari, B. Ringham-Terry, J. R. Wainwright, G. G. Jones, F. McCormick, P. Rodriguez-Viciana, SHOC2-MRAS-PP1 complex positively regulates RAF activity and contributes to Noonan syndrome pathogenesis. Proc Natl Acad Sci U S A. 115, E10576– E10585 (2018).

51. S. Dharmaiah, T. H. Tran, S. Messing, C. Agamasu, W. K. Gillette, W. Yan, T. Waybright, P. Alexander, D. Esposito, D. V. Nissley, F. McCormick, A. G. Stephen, D. K. Simanshu, Structures of N-terminally processed KRAS provide insight into the role of N-acetylation. Sci Rep. 9, 10512 (2019).

52. T. Taylor, J.-P. Denson, D. Esposito, Optimizing Expression and Solubility of Proteins in E. coli Using Modified Media and Induction Parameters. Methods Mol Biol. 1586, 65–82 (2017).

53. F. W. Studier, Protein production by auto-induction in high density shaking cultures. Protein Expr Purif. 41, 207–234 (2005).

54. C. Agamasu, R. Ghirlando, T. Taylor, S. Messing, T. H. Tran, L. Bindu, M. Tonelli, D. V. Nissley, F. McCormick, A. G. Stephen, KRAS Prenylation Is Required for Bivalent Binding with Calmodulin in a Nucleotide-Independent Manner. Biophys J. 116, 1049–1063 (2019).

55. K. Kopra, E. Vuorinen, M. Abreu-Blanco, Q. Wang, V. Eskonen, W. Gillette, A. T. Pulliainen, M. Holderfield, H. Härmä, Homogeneous Dual-Parametric-Coupled Assay for Simultaneous Nucleotide Exchange and KRAS/RAF-RBD Interaction Monitoring. Anal Chem. 92, 4971–4979 (2020).

56. T. Travers, C. A. López, C. Agamasu, J. J. Hettige, S. Messing, A. E. García, A. G. Stephen, S. Gnanakaran, Anionic Lipids Impact RAS-Binding Site Accessibility and Membrane Binding Affinity of CRAF RBD-CRD. Biophys J. 119, 525–538 (2020).

57. F. Delaglio, S. Grzesiek, G. W. Vuister, G. Zhu, J. Pfeifer, A. Bax, NMRPipe: a multidimensional spectral processing system based on UNIX pipes. J Biomol NMR. 6, 277–293 (1995).

58. W. Lee, M. Tonelli, J. L. Markley, NMRFAM-SPARKY: enhanced software for biomolecular NMR spectroscopy. Bioinformatics. 31, 1325–1327 (2015).

59. D. W. Morgens, M. Wainberg, E. A. Boyle, O. Ursu, C. L. Araya, C. K. Tsui, M. S. Haney, G. T. Hess, K. Han, E. E. Jeng, A. Li, M. P. Snyder, W. J. Greenleaf, A. Kundaje, M. C. Bassik, Genome-scale measurement of off-target activity using Cas9 toxicity in high-throughput screens. Nat Commun. 8, 15178 (2017).

60. K. L. Schreiber, L. Paquet, B. G. Allen, H. Rindt, Protein kinase C isoform expression and activity in the mouse heart. Am J Physiol Heart Circ Physiol. 281, H2062–2071 (2001).

61. J. Schindelin, I. Arganda-Carreras, E. Frise, V. Kaynig, M. Longair, T. Pietzsch, S. Preibisch, C. Rueden, S. Saalfeld, B. Schmid, J.-Y. Tinevez, D. J. White, V. Hartenstein, K. Eliceiri, P. Tomancak, A. Cardona, Fiji: an open-source platform for biological-image analysis. Nat Methods. 9, 676–682 (2012).

62. R. Edgar, M. Domrachev, A. E. Lash, Gene Expression Omnibus: NCBI gene expression and hybridization array data repository. Nucleic Acids Res. 30, 207–210 (2002).

